# Human iPSC 4R tauopathy model uncovers modifiers of tau propagation

**DOI:** 10.1101/2023.06.19.544278

**Authors:** Celeste Parra Bravo, Alice Maria Giani, Jesus Madero Perez, Zeping Zhao, Avi Samelson, Man Ying Wong, Alessandro Evangelisti, Li Fan, Tatyana Pozner, Maria Mercedes, Pearly Ye, Tark Patel, Allan Yarahmady, Gillian Carling, Virginia M. Y. Lee, Manu Sharma, Sue-Ann Mok, Wenjie Luo, Mingrui Zhao, Martin Kampmann, Shiaoching Gong, Li Gan

## Abstract

Tauopathies are age-associated neurodegenerative diseases whose mechanistic underpinnings remain elusive, partially due to lack of appropriate human models. Current human induced pluripotent stem cell (hiPSC)-derived neurons express very low levels of 4-repeat (4R)-tau isoforms that are normally expressed in adult brain. Here, we engineered new iPSC lines to express 4R-tau and 4R-tau carrying the P301S *MAPT* mutation when differentiated into neurons. 4R-P301S neurons display progressive Tau inclusions upon seeding with Tau fibrils and recapitulate features of tauopathy phenotypes, including shared transcriptomic signatures, autophagic body accumulation, and impaired neuronal activity. A CRISPRi screen of genes associated with Tau pathobiology identified over 500 genetic modifiers of Tau-seeding-induced Tau propagation, including retromer VPS29 and the UFMylation cascade as top modifiers. In AD brains, the UFMylation cascade is altered in neurofibrillary-tangle-bearing neurons. Inhibiting the UFMylation cascade suppressed seeding-induced Tau propagation. This model provides a powerful platform to identify novel therapeutic strategies for 4R tauopathy.

## INTRODUCTION

Tauopathies, characterized by accumulation of tau aggregates, are a heterogeneous group of neurodegenerative diseases. They include Alzheimer’s disease (AD), the most common tauopathy, as well as frontotemporal lobar degeneration with Tau pathology (FTLD-Tau), corticobasal degeneration (CBD), progressive supranuclear palsy (PSP), argyrophilic grain disease (AGD), globular glial tauopathy, chronic traumatic encephalopathy (CTE), and Pick’s disease (PiD) (Gotz et al., 2019). The microtubule-associated protein Tau is encoded by a single gene (*MAPT*) and gives rise to six isoforms, including isoforms containing either three (3R) or four (4R) microtubule-binding repeats, due to alternative splicing of exon 10 (Goedert et al., 1989). Based on the dominant 3R or 4R isoforms, there are three subtypes of tauopathies, 3R, 4R, and 3R/4R mixed tauopathies that exhibit distinct Tau filament structures revealed by cryogenic electron microscopy (cryo-EM). Tau filaments in AD (3R/4R) (Fitzpatrick et al., 2017), PiD (3R), CBD (4R), and PSP (4R) (Shi et al., 2021) are structurally distinct. Among the *MAPT* mutations that cause familial cases of FTLD-tau, many alter the ratio of 3R to 4R (Hutton et al., 1998; Spillantini et al., 1998), and several of the mutations, including P301S/L (Mirra et al., 1999), are located in the exon 10 and therefore 4R-specific.

Human iPSC-derived neurons, especially those derived from mutation-carrier patients, are invaluable in modeling neurological diseases, including tauopathies (Karch et al., 2019; Paonessa et al., 2019). Combined with CRISPR-Cas9 technology, iPSC-derived neuronal platforms enable isogenic controls for precise disease modeling (Sohn et al., 2019) and functional genomics to identify disease modifiers (Tian et al., 2019). However, iPSC-derived neurons express very low levels of 4R-Tau even after extended periods of culture and are thus unsuitable to model 4R tauopathy, such as PSP (Sposito et al., 2015; Verheyen et al., 2015). The low levels of exon-10-containing tau also limit their relevance in modeling the dominant familial FTLD-tau mutations located in exon 10 (Paonessa *et al*., 2019). Moreover, it has been difficult to capitulate robust Tau aggregation in human iPSC-derived neurons. While no insoluble Tau aggregates were observed in MAPT-P301L or MAPT-IVS10+16 iPSC neurons (Paonessa *et al*., 2019), limited Tau inclusions were observed in the processes after 120 days. One likely contributing factor lies in modeling a condition that would require many years in aging neurons in young iPSC-neurons in culture for weeks, while another important factor is the lack of 4R Tau in iPSC-derived neurons (Capano et al., 2022). In the current study, we report the establishment of a robust and scalable human iPSC 4R tauopathy model. We engineered new lines of iPSC-neurons to express 4R-tau and 4R-tau carrying the P301S *MAPT* mutation. Upon seeding with Tau fibrils, we showed that 4R-P301S neurons develop a progressive spread of Tau aggregation, aberrant neuronal activity, and endolysosomal pathway dysfunction. Using CRISPRi-based functional genomic screening, we identified novel genetic modifiers and pathways and established a robust platform to discover potential therapeutic strategies.

## RESULTS

### Generation and characterization of 4R-tau and human iPSC-derived neurons

Among the six isoforms of *MAPT*, the 4-repeat (4R) Tau, resulting from the inclusion of exon 10 via alternative splicing, expresses four microtubule-binding domains (MTBD) and plays important roles in the pathogenesis of tauopathies. However, modeling 4R tauopathy in iPSC neurons has been difficult since human iPSC-derived neurons, including i^3^Neurons (i^3^N) that express inducible Neurogenin-2 transcription factor, express very low levels of 4-repeat Tau (Sposito *et al*., 2015; Verheyen *et al*., 2015; Wang et al., 2017). To elevate 4R-tau expression, we edited the *MAPT* locus of i^3^Neurons via CRISPR/Cas9 using a donor plasmid containing point mutations at the 3′ and 5′ ends of exon 10 that prevent snRNP binding and splicing of the pre-mRNA (Figure 1A). After puromycin selection and FLP recombinase treatment to excise the Puro-GFP cassette, clones with one or both alleles of *MAPT* edited to express 4R-Tau were selected. Two 4R-homozygotic clones (clone #1 and clone #2) and one heterozygotic clone were confirmed with normal karyotypes and markers of pluripotency, including SOX2, SSEA4, NANOG, TRA-1-81, OCT3/4, and TRA-1-60 (Figure S1A-D).

**Figure 1.**
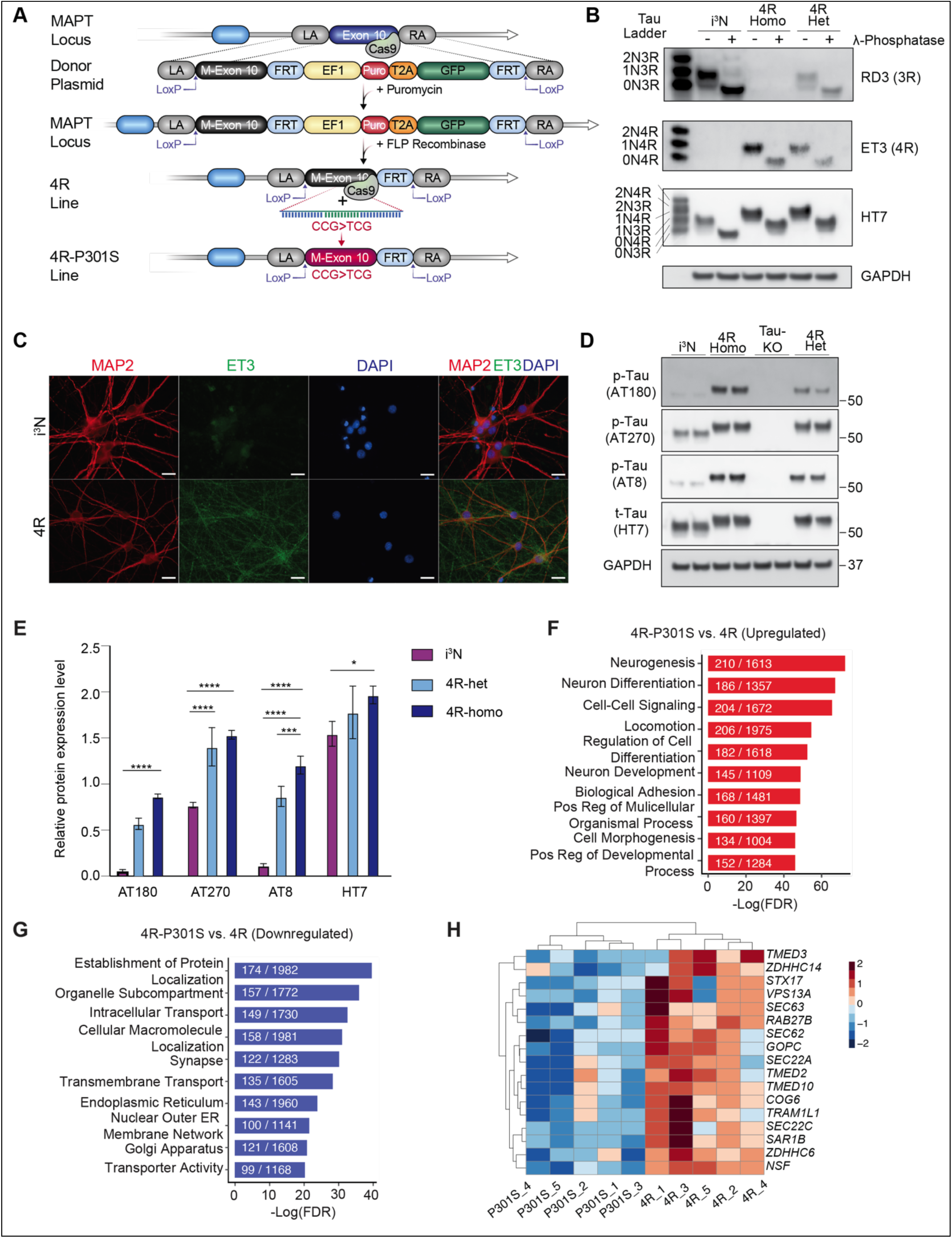
Generation and characterization of 4R-Tau human iPSC-derived neurons. (A) Stepwise strategy of CRISPR/Cas9-mediated genome editing to generate *MAPT* 4R and 4R-P301S knock-in. (B) Representative immunoblot of lysates from 3R (i^3^N), 4R heterozygous, and 4R homozygous 6-weekold neurons (D42) before and after lambda protein phosphatase treatment. n=2 independent experiments. (C) Representative immunofluorescence images of 6-week-old (D44) i3N and 4R neurons stained with MAP2, ET3, and DAPI. Scale bar, 25 μm. (D and E) Representative immunoblots (D) and the quantification (E) of phosphorylated tau (AT180, AT270, AT8) and total tau (HT7) in 6-week-old (D42) i^3^N, 4R homozygous, Tau-KO, and 4R homozygous neurons. Normalized to GAPDH. * p < 0.05, *** p < 0.001, **** p < 0.0001, one-way ANOVA, Tukey’s multiple comparisons test. (F and G) Gene set enrichment analysis pathways identified for upregulated (F) and downregulated (G) DEGs in 4R-P301S vs 4R neurons. Significantly enriched Gene Ontology terms for biological process, cellular component, and molecular function are shown. (H) Heatmap and hierarchical clustering of DE genes within the Transmembrane Transport and Intracellular Transport pathways in untreated 4R-P301S and 4R neurons. Upregulated genes > 0 (red), Downregulated genes < 0 (blue). The genes listed include DE genes based on read count, adjusted pvalue < 0.05. Gene Ontology term for biological process. Abbreviations: phosphorylated Tau (p-Tau), total Tau (t-Tau), 3R homozygous (i^3^N), 4R homozygous (4R).

We differentiated homozygotic clone #1 into excitatory neurons with the addition of doxycycline, as described (Wang *et al*., 2017). The parental i^3^N line and a heterozygotic line with one allele of *MAPT* locus edited were included as controls. Western blot analyses were performed using antibodies recognizing 3R MTBDs and ET3 specific for exon 10 (Espinoza et al., 2008), and HT7, a pan-tau antibody. Treatment with phosphatase allowed more accurate alignment with the six isoforms of recombinant Tau and served as molecular weight (MW) controls (Figure 1B). Compared to i^3^N neurons, which express only 3R-tau labeled with RD3 antibody, neurons derived from homozygotic 4R tau (clone #1) express exclusively 4R-tau labeled with ET3 (Figure 1B). In contrast, neurons derived from a normal karyotype 4R-hetero clone #1 (Figure S1C) express both 3R-tau and 4R-tau (Figure 1B). Immunocytochemistry confirmed the immunoreactivity of ET3, a 4R-specific antibody in 4R homozygous clone #1 neurons (Figure 1C) and clone #2 (Figure S1E), but not in i^3^N neurons (Figure 1C). Interestingly, compared with i^3^N neurons, both 4R homozygous and heterozygous neurons express higher levels of phospho-tau species, which have been associated with AD and primary tauopathies (Figure 1D, 1E).

One of the most common frontotemporal dementia (FTD)-linked *MAPT* mutations, P301S, is located in exon 10 (Yasuda et al., 2005). P301S is also highly aggregation-prone (Allen et al., 2002; Berriman et al., 2003). To model FTD tauopathy, we further edited the homozygous 4R-tau (4R) line with CRISPR/Cas9 to include the P301S mutation using a single-stranded DNA oligonucleotide and replacing a proline with a serine at residue 301 (Figure 1A, Figure S2A). Two clones of homozygous 4R-P301S-tau-expressing (4R-P301S) iPSCs (Figure S2B, S2C) were confirmed for normal karyotypes and pluripotency using markers (i.e., SOX2, SSEA4, NANOG, TRA-1-81, OCT3/4, TRA-1-60) (Figure S2D). To dissect the molecular alterations induced by the P301S mutation, we performed bulk RNA-sequencing analyses of 4R and 4R-P301S neurons (Figure S2). Unsupervised cluster and principal component analyses (PCA) revealed that 4R neurons and 4R-P301S neurons were clustered separately (Figure S2E-F), consistent with the pathogenic nature of P301S mutation. Over 2200 genes were altered by P301S mutation (Table S2). The upregulated genes showed enrichment in *Neuron differentiation* and *Cell-Cell signaling* pathways (Figure 1F), and the downregulated ones were enriched in genes related to *Organelle localization*, *Intracellular transport, and Transmembrane transport* (Figure 1G), some of which were highlighted in the heatmap (Figure 1H). The P301S mutation downregulated expression of Transmembrane p24 trafficking proteins (*TMED)*3, *TMED10,* and *TMED2* (Figure 1H), which are all involved in the transport of proteins between the ER and the Golgi apparatus and play essential roles in maintaining cellular homeostasis and proper folding and transport of proteins in the cell. Other downregulated genes include *Syntaxin-17* (*STX-17*), involved in the fusion of autophagosomes with lysosomes, and *Vacuolar Protein Sorting 13 Homolog A* (*VPS13A)*, an essential protein facilitating non-vesicular lipid transfer between organelles, such as the endoplasmic reticulum (ER) and mitochondria, or the ER and endosomes/lysosomes (Figure 1H). These findings suggest that 4R-P301S neurons are likely to be more vulnerable to proteostasis imbalance.

### Modeling propagation of 4R-tau inclusions

Using the MC1 antibody, a conformation-specific antibody that recognizes disease-specific forms of Tau from human patients (Jicha et al., 1997), we detected no obvious insoluble Tau aggregates in 4R and 4R-P301S neurons even after weeks in culture (Figure S3A). To model seeding-induced Tau propagation, we treated the 4R and 4R-P301S neurons with K18-P301L-tau (K18) fibrils, a truncated form of human Tau containing only the aggregation-prone repeat domain of the microtubule-binding domain (Gustke et al., 1994) (Figure 2A). Following a treatment period of 3–5-weeks, we detected robust MC1-positive inclusions only in 4R-P301S, not in 4R neurons (Figure 2B). Since MC1 immunoreactivity requires domains outside of K18, the MC1+ inclusions are made of endogenous 4R Tau upon seeding with Tau fibrils, a process reflecting templating-induced propagation of misfolded protein. Weekly analyses at 1–5 weeks post-seeding showed a progressive increase in MC1+ inclusions (Figure 2C, 2D), supporting a prion-like model of Tau propagation. Further analyses revealed that the observed MC1+ inclusions were immunoreactive to antibodies against oligomers and phosphorylation (Figure 2E). We also performed transmission electron microscopy (TEM) and detected prominent fibrillar Tau structures in the soma of 4R-P301S neurons seeded with tau (Figure 2F).

**Figure 2.**
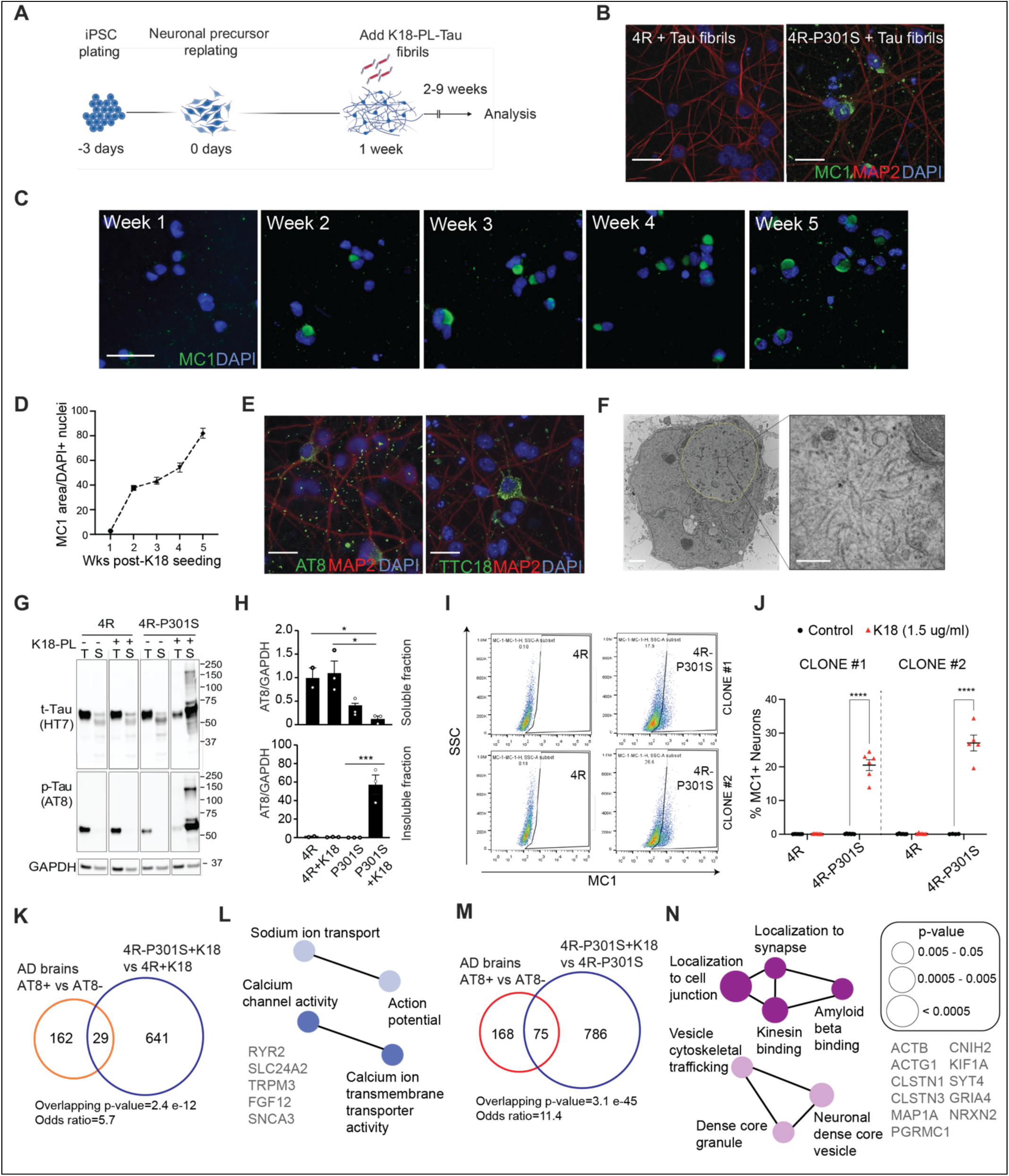
Modeling seeding-induced 4R-Tau inclusions in human neurons. (A) Diagram illustrating the two-step differentiation of 4R and 4R-P301S human iPSCs into doxycycline-inducible glutamatergic neurons, followed by K18-P301L Tau fibril (K18) seeding at 1 week, and used for experiments ≥2 weeks post-seeding. This figure was created with Biorender.com (B) Representative immunofluorescence images of K18-seeded D25(7+18) 4R and 4R-P301S neurons stained with MC1, MAP2, and DAPI. Scale bar, 25 μm. n=3, three independent experiments. (C and D) Representative images (C) and quantification (D) of immunofluorescence time course quantification of MC1/DAPI+ nuclei in 4R-P301S neurons 1–5 weeks post-K18 treatment. n=4, two independent experiments performed in replicate. Scale bar, 50 μm. (E) Representative immunofluorescence images of D25(7+18) 4R-P301S neurons seeded with K18 showing the presence of phosphorylated Tau (AT8), oligomeric Tau (TTC18), MAP2, and DAPI. Scale bar, 25 μm. (F) Left panel: Representative TEM image of a 3 μg/ml K18-seeded D35(7+28) 4R-P301S neuronal soma with a large tau inclusion (outlined). Scale bar, 2 μm. Right panel: Enlarged image showing distinct individual tau fibrils resolved upon magnification. Scale bar, 200 nm. (G and H) Representative immunoblot images (G) of detergent-fractionated lysates from D59(D7+52) 4R and 4R-P301S neurons -/+ 1.5 µg/ml K18 seeding stained with total tau (HT7) and p-Tau (AT8). T=Triton soluble, S=SDS soluble. Quantification (H) of lysates from D68(22+46) and D59(D7+52) 4R and 4R-P301S normalized to GAPDH. n=3, two independent experiments, one performed in duplicate. * p < 0.05, *** p < 0.001, one-way ANOVA, Tukey post-hoc test. (I and J) Representative flow cytometry analysis (I) and quantification (J) of the percentage of MC1+ cells in and D42(7+35) 4R and 4R-P301S neurons clone #1 and clone #2 -/+ 1.5 µg/mL K18 groups. n=3–6 biological replicates. **** p < 0.0001, two-way ANOVA, Šídák’s multiple comparisons test. (K) Venn diagram comparing the number and overlap of downregulated DEGs in 4R-P301S+K18 vs 4R+K18 neurons from bulk RNA-seq analysis, and AT8+ vs AT8-neurons from pseudo bulk RNA-seq analysis of AD brains. Fisher’s exact test. (L) ClueGO biological processes pathway enrichment of overlapping downregulated DEGs in 4R-P301S+K18 vs 4R+K18 / AT8+ vs AT8-comparison. Node colors denote functionally grouped networks (kappa connectivity score ≥ 40%). GO Term ≥ 1% of genes. p-value = 0.01-0.04. (M) Venn diagram comparing the number and overlap of upregulated DEGs in 4R-P301S+K18 vs 4R-P301S neurons from pseudo bulk RNA-seq analysis, and AT8+ vs AT8-neurons from pseudo bulk RNA-seq analysis of AD brains. Fisher’s exact test. (N) ClueGO biological processes pathway enrichment of overlapping upregulated DEGs in 4R-P301S+K18 vs P301S / AT8+ vs AT8-comparison. Node colors denote functionally grouped networks (kappa connectivity score ≥ 40%). GO Term ≥ 4% of genes.

To further analyze the biochemical nature of the inclusions, lysates were first solubilized by Triton-X, and the insoluble fractions were further solubilized by SDS. In 4R neurons, the majority of Tau and phospho-Tau was observed in the Triton-soluble fraction with no high-molecular-weight (MW) Tau and phospho-Tau observed, consistent with the lack of MC1+ aggregates with or without K18 Tau seeding (Figure 2G). Quantification confirmed that Tau-seeding of 4R-P301S neurons significantly reduced levels of AT8-positive phospho-Tau in the Triton-soluble fraction, while markedly elevating those in the Triton-insoluble fraction (Figure 2H). Another species of phospho-tau (AT270) was also observed in the Triton-insoluble/SDS fraction of 4R-P301S neurons seeded with K18-Tau fibrils (Figure S3B).

Next, we used flow cytometry with MC1 antibody to measure somatic Tau inclusions quantitatively. We examined the extent of Tau inclusions at two times, using two independent 4R-P301S clones to confirm the reproducibility of the phenotype. At Day 21 post-seeding of K18 fibrils (1.5 µg/ml), both clone #1 and clone #2 4R-P301S neurons exhibited highly consistent levels of MC1+ population of neurons (Figure S3C, S3D). The percentage of MC1+ neurons increased 5–6-fold between Day 21 and Day 42 post-seeding in both clone #1 and clone #2, reflecting seeding-induced propagation/amplification (Figure 2I, 2J). Increasing the Tau fibrils from 1.5 µg/ml to 3.0 µg/ml resulted in a modest increase in Tau inclusions in both 4R-P301S clones, either at Day 21 (Figure S3E, S3F) or Day 42 (Figure S3G, S3H) post-seeding. To address whether the mutant tau fibrils induce species-specific templating of mutant tau in 4R-P301S neurons, we seeded neurons with WT 0N4R-tau fibrils for 3 weeks and detected MC1-positive inclusions only in 4R-P301S, not in 4R neurons (Figure S3I). Thus, Tau aggregation in 4R-P301S neurons is not limited to seeding with fibrils with the same mutated residue.

Molecular signatures of tangle-bearing neurons in AD were recently characterized by single-cell RNA sequencing (Otero-Garcia et al., 2022). To interrogate the transcriptomic alterations in MC1+ Tau inclusion-bearing neurons, we performed bulk RNA-seq and identified the differentially expressed genes (DEGs) in 4R-P301S+K18 that developed seeding-induced inclusions and 4R+K18 neurons that did not have inclusions, despite the addition of K18 seeds (Figure S4A, S4B, Table S3). To assess gene expression similarities between our *in vitro* platform and post-mortem human AD brains, we performed a gene overlap analysis of the DEGs with those comparing AT8+ vs AT8-excitatory neurons in human AD brains (Otero-Garcia *et al*., 2022) (Figure 2K). A modest yet significant gene overlap was detected in the downregulated DEGs (odds ratio = 5.7, p-value = 2.4e-12). Gene Ontology (GO) enrichment analyses revealed that the molecular functions of these overlapping proteins largely comprised action potential and calcium transport networks (Figure 2L). To exclude possible effects of the P301S mutation, we next performed single-cell RNA sequencing to compare the transcriptomes of tau-seeded vs. non-seeded 4R-P301S neurons (Figure S4C-E). We reasoned that the pseudo-bulk analyses of the DEGs would be enriched with molecular signatures of inclusion-containing human neurons (Table S4). Indeed, comparison with AT8+ vs AT8-excitatory neurons in human AD brains revealed a highly significant overlap in the upregulated DEGs (odds ratio = 11.4, p-value = 3.1e-45, Figure 2M). Synaptic localization and vesicular trafficking networks are some of the prominent overlapping pathways (Figure 2N).

### Endolysosomal dysfunction in 4R-P301S neurons promotes Tau propagation

4R-P301S neurons exhibited striking downregulated pathways in *Organelle localization*, *Intracellular transport, and Transmembrane transport* (Figure 1G, 1H). To further dissect alterations of cellular machinery caused by Tau inclusions, we next performed TEM analyses to compare subcellular structures in 4R and 4R-P301S neurons with or without K18 seeding. Marked accumulation of abnormal vesicular structures with multilayer membranes was observed exclusively in 4R-P301S neurons treated with K18 fibrils, not in 4R neurons or 4R-P301S neurons without seeding (Figure 3A, 3B). These abnormal structures were reminiscent of multilamellar bodies (MLBs), organelles containing multiple concentric membrane layers of lysosomal origin (Hariri et al., 2000). MLBs were observed in the soma along with Tau inclusions (Figure 3C), as well as in the processes of 4R-P301S neurons (Figure 3D).

**Figure 3.**
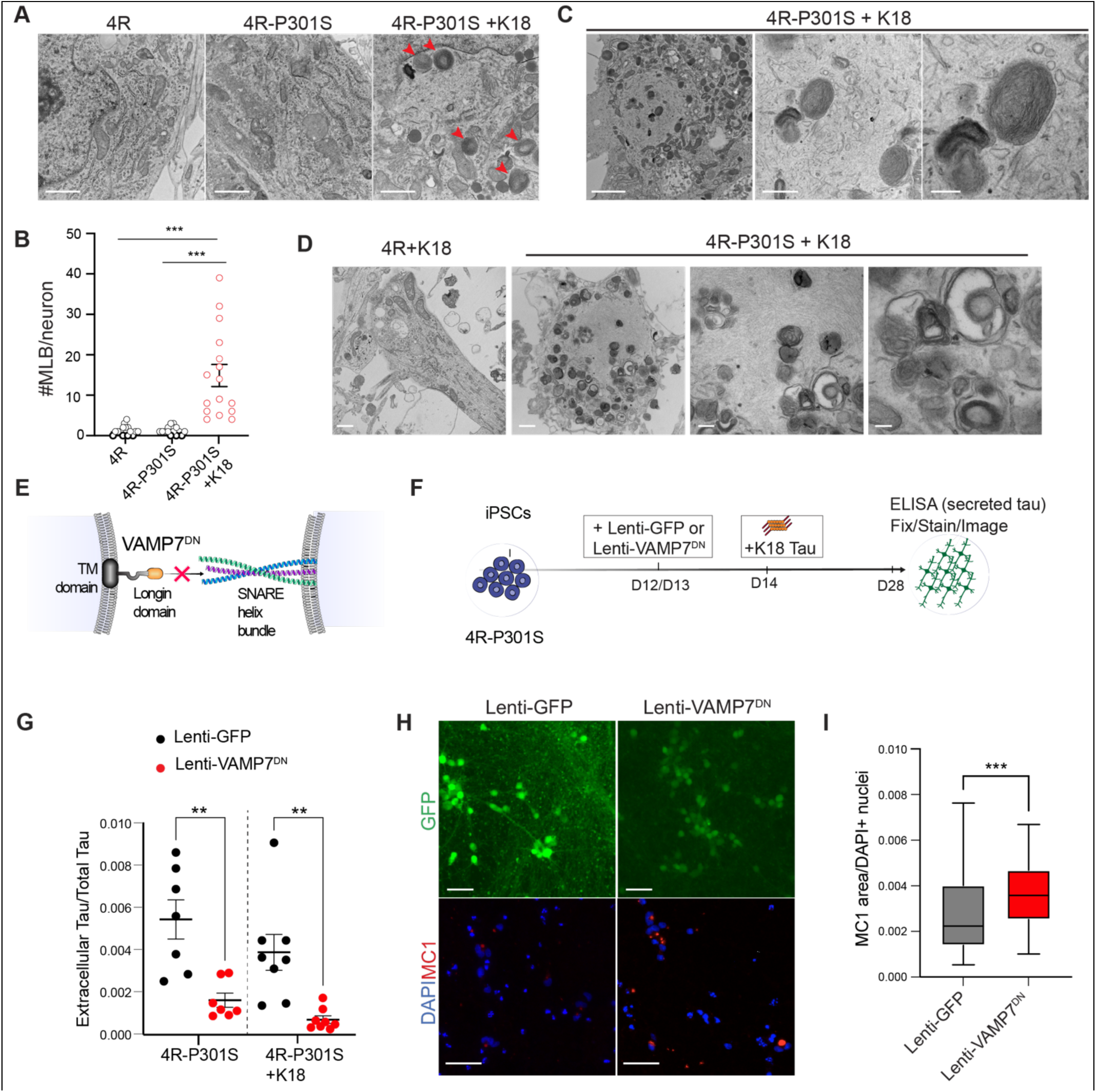
Endolysosomal dysfunction promotes propagation of Tau inclusions. (A) Representative TEM images from soma D35 4R neurons, D40 4R-P301S neurons, and 1.5 µg/mL K18-seeded D40(7+33) 4R-P301S neurons. Red arrows = MLB. Scale bar, 1 μm. (B) Quantification of #MLB/neuron from D35 and D40 4R and 4R-P301S, and D43(7+36) 4R-P301S+K18 neurons (3 µg/mL) from TEM images. n=4, 2 replicates per group, two independent experiments. *** p < 0.001, one-way ANOVA, Tukey post-hoc test. (C) Representative TEM images from soma of 3 µg/mL K18-seeded D43(7+36) 4R-P301S neurons at different magnifications. Scale bars left to right: 2 μm, 500 nm, 200 nm. (D) Representative TEM images from neuronal processes of 3 µg/mL K18-seeded D35(7+28) 4R and 4R-P301S neurons showing dystrophic neurites presenting with Tau inclusions and increased numbers of MLBs in 4R-P301S neurons. Scale bars left to right: 500, 500, 200, and 100 nm.

The accumulation of MLBs in 4R-P301S neurons with Tau inclusions strongly suggests dysfunctional endolysosomal membrane trafficking. Since lysosomal exocytosis has been linked with the spread of misfolded protein aggregates, including alpha-synuclein (Xie et al., 2022), we reasoned that impaired lysosomal fusion could be involved in our seeding-induced tauopathy model. To block lysosomal fusion, we inhibited the activity of VAMP7, a calcium-dependent v-SNARE protein that mediates lysosomal membrane fusion (Arantes and Andrews, 2006), by overexpressing dominant-negative VAMP7 (VAMP7^DN^) (Xie *et al*., 2022) (Figure 3E). 4R-P301S neurons were infected with Lenti-GFP-VAMP7^DN^ or Lenti-GFP-control, followed by seeding with K18 fibrils, and analyses were performed 2 weeks later (Figure 3F). Perturbing VAMP7-mediated membrane fusion significantly reduced the amount of extracellular Tau released after KCl-induced depolarization, relative to total Tau, measured with ELISA as described (Tracy et al., 2022) (Figure 3G). Strikingly, blockage of VAMP7 activity resulted in a significant increase in MC1+ Tau inclusions normalized to cell number (Figure 3H, 3I). Together, our results suggest that impaired membrane fusion, possibly lysosomal exocytosis, exacerbates seeding-induced Tau propagation.

### Tau inclusions impair neuronal activity

Tau pathology correlates strongly with cognitive decline (Bejanin et al., 2017; Love et al., 2014; Ossenkoppele et al., 2022). However, how neuronal functions are affected by Tau inclusions remains poorly defined due to the challenge of labeling Tau inclusions in live neurons. To track live human neurons with or without Tau aggregates for calcium imaging, we modified the 4R-P301S line to insert a HaloTag at the 5′ end of the *MAPT* locus using CRISPR-Cas9 gene editing (Figure 4A). Upon the binding of the HaloTag to a synthetic ligand of choice, HaloTag undergoes a conformational change and fluoresces at a given wavelength, enabling live-cell imaging of endogenous Tau molecules (Grimm et al., 2015). Two independent homozygous 4R-P301S-HaloTag iPSC clones were confirmed for normal karyotypes (Figure S5A-C), and clone #1 was used for functional experiments. Upon addition of JFX-549 synthetic ligand (Grimm *et al*., 2015), 4R-P301S neurons with and without obvious aggregated Tau can be visually distinguished at 568 nm (Figure 4B).

**Figure 4.**
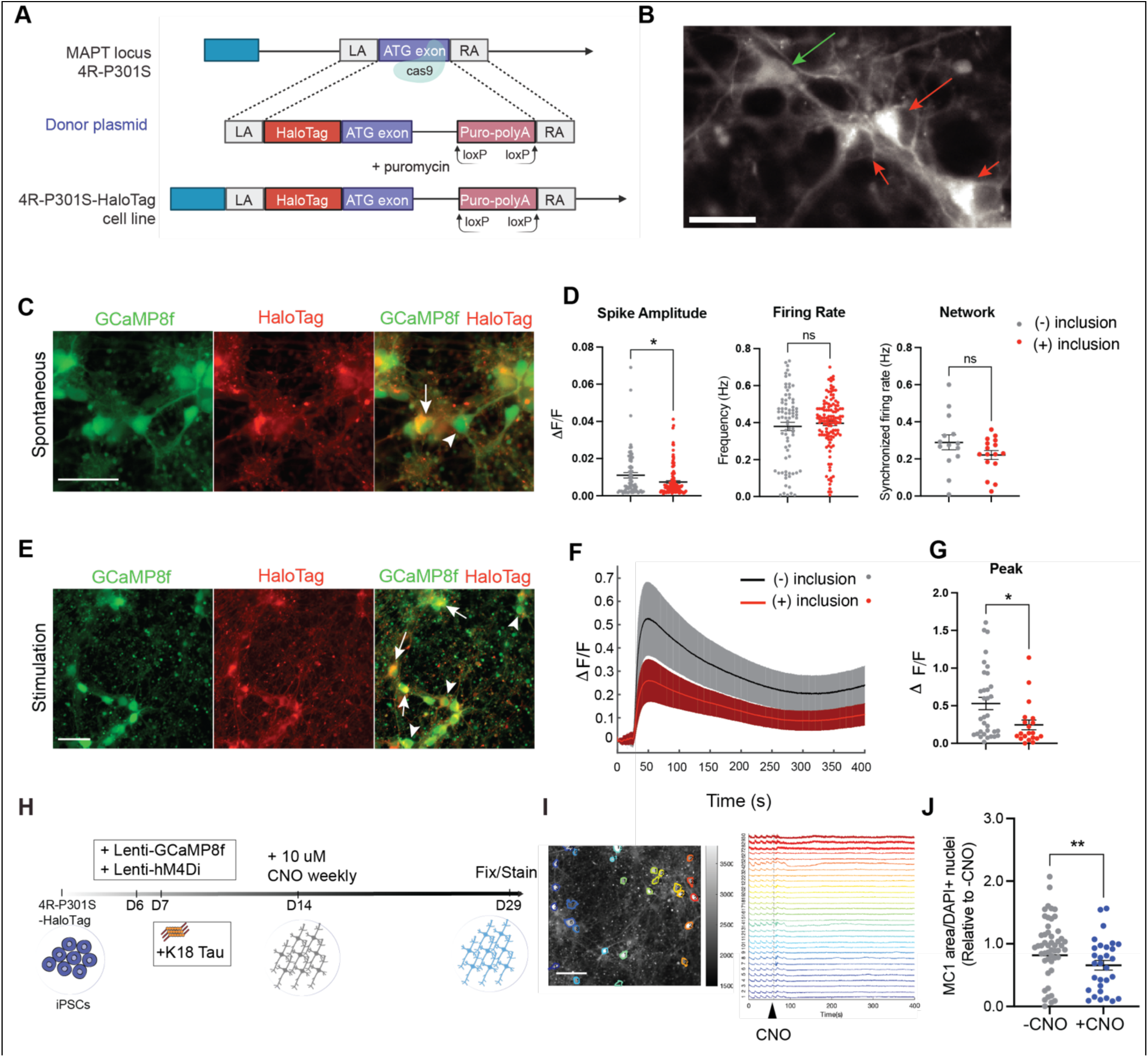
Tau inclusions impair spontaneous and evoked neuronal activity. (A) Schematic illustrating the CRISPR-mediated knock-in of HaloTag cassette at 5′ end of *MAPT* locus in the 4R-P301S iPSC line. This figure was created with Biorender.com (B) Representative fluorescence image of 3 µg/ml K18-seeded 4R-P301S-HaloTag neurons (D24) harboring Tau inclusions (green arrow: *-inclusion*; red arrow: *+inclusion*). Incubated with JFX_549_-HaloTag ligand. Scale bar, 50 μm. (C) Representative fluorescence images expressing GCaMP8f, HaloTag, and merged two sensors in D24 K18-seeded 4R-P301S-HaloTag spontaneous activity. white arrowhead: *-inclusion*; white arrow: *+inclusion*. Scale bar, 50 μm. (D) Quantification of spike amplitude, firing rate, and synchrony index from calcium imaging of K18-seeded 4R-P301S-HaloTag neurons (D24-30). n=3 replicates, two independent experiments. *p < 0.05, unpaired t-test. (E) Representative fluorescence images of GCaMP8f, HaloTag, and merged two sensors in D30 K18-seeded 4R-P301S-HaloTag treated with 50 mM KCl. white arrowhead: *-inclusion examples*; white arrow: *+inclusion examples*. Scale bar, 50 μm.

To monitor neuronal activity simultaneously, we transduced seeded 4R-P301S neurons with a new generation of genetically encoded calcium sensor GCaMP8-fast (Lenti-hSynapsin-hGCaMP8f) (Zhang et al., 2023) (Figure 4C, S5D). Calcium transients, measured by the change in fluorescence over basal fluorescence (dF/F), were recorded over 2 minutes per somal region of interest (ROI) within an image frame (Figure 4C). ROIs in the soma were then classified into visually distinguishable *-inclusion* or *+inclusion* bins, and spontaneous neuronal activities from single neurons were extracted from the calcium trace (Figure 4C). Compared with neurons lacking inclusions, those with inclusions displayed a significant decrease in spike amplitudes, whereas the firing and synchronicity rates remained unchanged (Figure 4D). To measure evoke responses, 50 mM KCl was perfused to induce chronic depolarization (Figure 4E). Neurons with visible somatic inclusions exhibited significant lower trace peak heights than those without visible inclusions (Figure 4F, 4G). Thus, both spontaneous and stimulation-evoked neuronal activity were impaired in human neurons with Tau inclusions.

### Neuronal activity promotes Tau propagation

The release of Tau is enhanced by neuronal activity in mouse brains and human neurons (Pooler et al., 2013; Tracy *et al*., 2022; Wu et al., 2016; Yamada et al., 2014). FTD mutations, such as V337M, induce hyperexcitability in human neurons (Sohn *et al*., 2019). To directly examine the effects of neuronal activity on Tau seeding and spread, we induced chronic silencing of neuronal activity by transducing the viral vector encoding an inhibitory DREADD receptor, hM4Di, genetically modified to respond specifically to the synthetic ligand clozapine-N-oxide (CNO), resulting in suppression of neuronal firing (Zhu and Roth, 2014). 4R-P301S-HaloTag neurons were infected with mCherry-hM4Di lentivirus, followed by K18 seeding (Figure 4H). One week after the seeding, CNO was applied to the culture weekly for 2 additional weeks to determine the outcome of chronic suppression of neuronal activity (Figure 4H). Robust expression of the mCherry-hM4Di receptor was confirmed before and after CNO administration (Figure S5E). Using calcium imaging, we confirmed that neuronal activities were silenced upon addition of CNO (Figure 4I). Levels of Tau inclusions, measured with MC1+ aggregates, were significantly reduced by CNO treatment compared to those with vehicle treatment (Figure 4J). Thus, neuronal activity promotes seeding-induced Tau propagation in human neurons.

### CRISPRi screen identifies genetic modifiers for tau propagation in 4R tauopathy

This new model of 4R Tau propagation is engineered from our i^3^N neuron platform, which facilitates the large-scale production of iPSC-derived glutamatergic neurons and enables scalable CRISPRi-based functional genomics (Tian et al., 2019). To identify genetic modifiers of Tau propagation, we engineered 4R-P301S iPSCs to stably express dCas9 by inserting CAG promoter-driven dCas9-BFP-KRAB into the CLYBL safe harbor locus by TALENs (Tian *et al*., 2019). After FACS sorting for BFP and selection with puromycin, two clones of heterozygous 4R-P301S-dCas9 lines were selected and confirmed for normal karyotype, clone #1 (Figure S6A) and clone #2 (Figure S6B). 4R-P301S-dCas9 iPSC clone #1 was stained for pluripotency markers OCT4, TRA-1-60, NANOG, TRA-1-81, SOX2, SSEA4 (Figure S6C) and was used for subsequent CRISPRi experiments. The knockdown efficiency of the 4R-P301S-dCas9 iPSC clone #1 was confirmed by TFRC staining and flow cytometry (Figure S6D, S6E).

The 4R-P301S-dCas9 iPSCs were transduced with a custom lentiviral CRISPRi sgRNA library targeting genes involved in Tau pathobiology based on a genome-wide CRISPRi screen for modifiers of tau oligomer levels in human iPSC-derived neurons (Samelson et al., 2023). This library consists of sgRNAs targeting 1,073 genes with five sgRNAs per gene and 250 non-targeting control (NTC) sgRNAs (Table S5). After library transduction, iPSCs were differentiated into neurons, and K18-Tau was seeded at D7 (Figure 5A). Neurons were collected at D19, fixed, and stained for MC1, and FACS sorted based on MC1 signal in the 488 nm channel into either MC1+ or MC1-bins (Figure 5A). Frequencies of cells expressing each sgRNA were determined by next-generation sequencing to uncover genes for which sgRNAs showed significant changes in frequency, either an increase or a decrease in relation to Tau inclusion phenotype (Figure 5A and Table S6).

**Figure 5.**
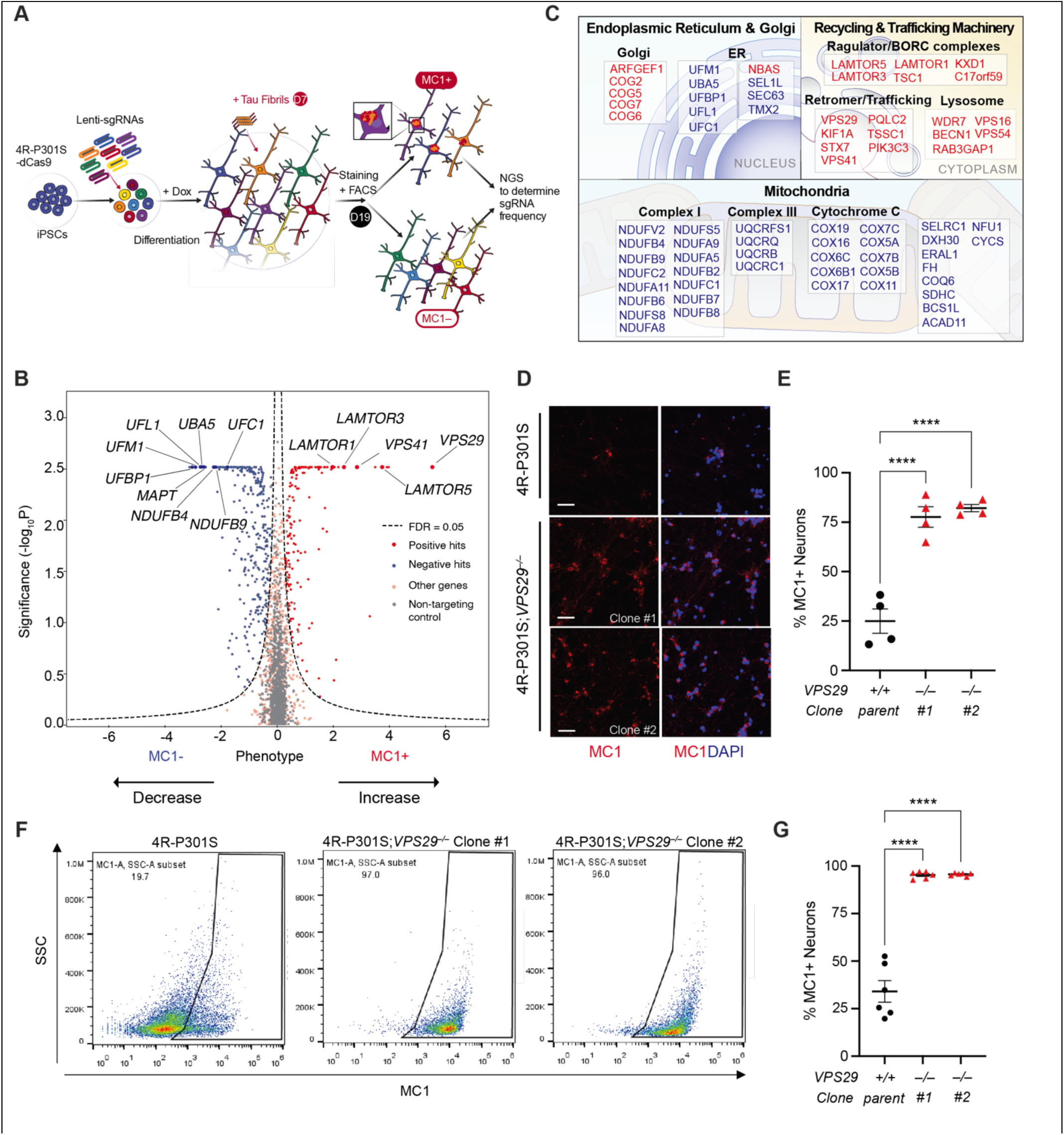
Identification of Tau inclusion modifiers by CRISPRi screening. (A) Strategy for CRISPRi screening based on Tau inclusions in 4R-P301S neurons. 4R-P301S iPSCs were transduced with pC13N-dCas9-BFP-KRAB lentivirus to express dCas9, generating 4R-P301S-dCas9 iPSCs. iPSCs were transduced with a custom sgRNA library lentivirus targeting genes involved in Tau aggregation (1073 genes) and differentiated, seeded with K18-Tau at D7, and collected at D19. Neurons were stained for MC1 and FACS-sorted to separate MC1+ and MC1-populations. Frequencies of neurons expressing a given sgRNA were determined by next-generation sequencing. (B) Volcano plot summarizing knockdown phenotypes and statistical significance (Mann-Whitney U test) for genes targeted in the pooled screen. MC1+, genes reducing Tau inclusions. MC1-, genes enhancing Tau inclusions. Dashed lines: cutoff for hit genes (FDR = 0.05).

As expected, *MAPT* sgRNAs were enriched in cells with reduced Tau inclusions (Figure 5B). Many of the other hits that reduced Tau inclusions are mitochondria genes, including many components in complex I *(NDUFB9*, *NDUFV2*, *NDUFB4*), complex II (*UQCRQ, UQCRB*), and cytochrome C (*COX6C, COX7B, COX7C*) (Figure 5B, 5C). We also observed enrichment of genes involved in UFMylation, a post-translational modification analogous to ubiquitylation (*UBA5, UFM1, UFBP1, UFL1, UFC1*) (Figure 5B, 5C) related to processes such as endoplasmic reticulum (ER)-associated protein degradation, ribosome-associated protein quality control at the ER, and ER-phagy (Eldeeb et al., 2021; Liang et al., 2020). Gene Ontology enrichment analyses revealed striking enrichment of distinct mitochondria and biosynthesis-related pathways (Figure S6G).

The top genes whose knockdown increased Tau inclusions were highly enriched in multiple pathways of vesicular trafficking, as revealed with Gene Ontology enrichment analyses (Figure S6F). Top hits include genes important for endolysosomal biogenesis and trafficking, such as LAMTOR complex *(LAMTOR1, LAMTOR3, LAMTOR5),* known as Late Endosomal/Lysosomal Adaptor and MAPK and mTOR Activator, as well as core subunits of HOPS (Homotypic fusion and Protein Sorting) complex, VPS16, and VPS41, which are critical for endosome-to-lysosome trafficking and fusion (Figure 5C). Genes involved in Golgi trafficking, such as Conserved Oligomeric Golgi (COG) complex (COG2, COG5, COG6, COG7) (Sumya et al., 2023) are also strong modifiers of Tau aggregation. Inhibition of VPS54, a subunit of the GARP (Golgi-Associated Retrograde Protein) complex, also exacerbated Tau inclusions (Figure 5B, 5C). VPS54 is involved in the retrograde transport of vesicles from endosomes to the trans-Golgi network (TGN). Notably, VPS29, a subunit of the retromer complex, emerged as one of the strongest modifiers. The retromer complex is responsible for the retrograde transport of cargo from endosomes to the Golgi apparatus and the plasma membrane, and VPS29 contributes to the cargo recognition and stability of the retromer complex.

To validate the hits from the CRISPRi screen, we deleted *VPS29* in 4R-P301S lines. Three independent clones of 4R-P301S;*VPS29*^−/−^ lines were generated and confirmed with normal karyotypes and markers of pluripotency (Figure S7A-C). Using qRT-PCR, we confirmed the deletion of *VPS29* in clones #1 and #2 (Figure S7D). All three clones and the 4R-P301S parental line were differentiated into neurons and seeded with K18-Tau fibrils. Three weeks after K18-Tau seeding, immunocytochemistry revealed a striking increase of MC1+ inclusions in all three clones of 4R-P301S;*VPS29*^−/−^, from 20% in the parental line to >80% in 4R-P301S;*VPS29*^−/−^ neurons (Figure 5D, 5E). To further confirm the effects of VPS29, we performed flow cytometry analyses in two of the clones. Strikingly, more than 90% of 4R-P301S;*VPS29*^−/−^ neurons exhibit MC1+ Tau inclusions, compared with 20–30% in the 4R-P301S neurons (Figure 5F, 5G), establishing VPS29 as a predominant modifier of Tau propagation in our 4R human tauopathy model.

### UFMylation cascade as a novel modifier of Tau propagation

We next selected the UFMylation cascade to validate the sgRNA hits that reduce Tau inclusions. Among the top hits are five out of six components in the UFMylation pathway. UBA5 and UFL1 are E1-like enzymes that activate UFM1, UFC1 is an E2-like enzyme that catalyzes activated UFM1 to target proteins, and UFBP1 is involved in the recognition and binding of UFMylated substrates (Figure 6A). Lentiviral constructs expressing UBA5, UFM1, or UFBP1 shRNAs under the U6 promoter and GFP under the SV40 promoter were generated for knockdown in human iPSC neurons. Using qRT-PCR, we confirmed an efficient knockdown of UBA5 and UFM1 (Figure S8C, S8D), but knockdown of UFBP1 resulted in significant toxicity (data not shown). In flow cytometry analyses performed 21 days post-seeding, the MC1+ signal was measured in GFP+ neurons, so only those expressing the given shRNA were included (Figure S8A). Seeding induced a robust MC1+ inclusions in neurons infected with control GFP-expressing lentivirus. However, MC1+/GFP+ neurons were markedly fewer with Lenti-shUBA5 or Lenti-shUFM1 than with Lenti-GFP controls (Figure 6B, 6C). MC1+/GFP+ neurons were also quantified using immunocytochemistry. In agreement with flow cytometry analyses, infection with Lenti-shUBA5 or Lenti-shUFM1 results in significantly fewer MC1+/GFP+ neurons than the Lenti-GFP controls (Figure 6D, 6E). The inclusion-promoting effects of UFMylation are not due to its effect on *MAPT* mRNA since levels of *MAPT* mRNA were not affected by Lenti-shUBA5 nor Lenti-shUFM1 (Figure S8B).

**Figure 6.**
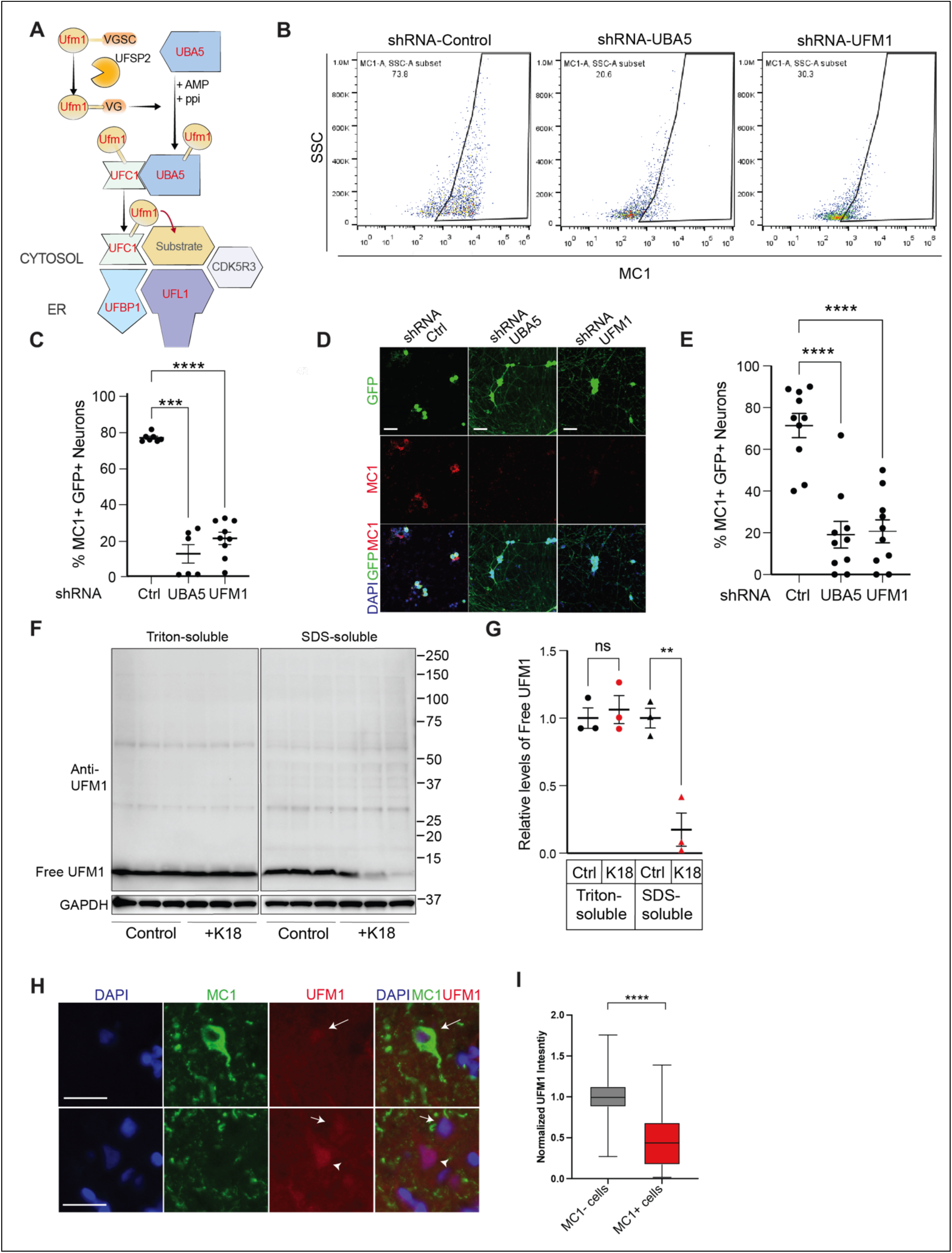
CRISPRi screen identifies the UFMylation pathway as a novel key modulator of Tau inclusions. (A) Diagram illustrating the protein components necessary for UFMylation of a target substrate. Abbreviation: VGSC, VGSC is an amino acid motif (valine-glycine-serine-cysteine). (B and C) Representative flow cytometry analysis (B) and the quantification (C) of the percentage of MC1+ cells in D21(7+14) 3 µg/ml K18-seeded 4R-P301S neurons infected with lentivirus (GFP+) containing shRNAs targeting UFMylation cascade genes (*UBA5, UFM1*). n=3-6 replicates, two independent experiments, ***p < 0.001, ****p < 0.0001, mixed model analysis, Dunnett’s multiple comparisons test. (D and E) Representative immunofluorescence images (D) and the quantification (E) of the percentage of DAPI+/MC1+ cells in D21(7+14) 3 µg/ml K18-seeded 4R-P301S neurons infected with lentivirus (GFP+) containing shRNAs targeting UFMylation cascade genes (*UBA5, UFM1*). n=6 replicates, two independent experiments. **** p < 0.0001, one-way ANOVA, Dunnett’s multiple comparisons test. Scale bar, 50 μm. (F) Representative immunoblot of relative free UFM1 D21 (7+14) 3 µg/ml K18-seeded 4R-P301S-HaloTag detergent-fractionated neuronal samples. Triton soluble (soluble), SDS-soluble (insoluble). Normalized to GAPDH. n=3 replicates per group, two independent experiments. (G) Quantification of relative free UFM1 in Triton-soluble and SDS-soluble fractions of control vs. K18-seeded neurons. ** p < 0.01, unpaired t-test. (H) Representative immunofluorescence images of human AD post-mortem tissue stained with DAPI, MC1, and UFM1. Arrow, MC1+ neuron. Arrowhead, MC1-neuron. Scale bar, 25 μm. (I) Quantification of UFM1 immunofluorescence normalized fold-change intensity in MC1+ and MC1-neurons in human AD post-mortem tissue. n=four male cases and three female cases, 12–15 neurons per group, per case. **** p < 0.0001, mixed model analysis.

We next compared the levels of UFMylation in K18-seeded and unseeded 4R-P301S neurons in Triton-soluble and Triton-insoluble (SDS-soluble) lysates, a fraction enriched with high MW Tau species from 4R-P301S neurons with Tau inclusions. Immunoblotting for an anti-UFM1 antibody preferentially recognizes free UFM1, revealed a strong reduction of free UFM1 (9.1 kDa) only in the triton-insoluble fraction of seeded 4R-P301S neurons (Figure 6F, 6G). To further explore the relevance of UFMylation in Tau pathology in AD, we performed co-immunostaining of MC1 and UFM1. In AD brains, we showed that MC1+ tangle-bearing neurons exhibited lower levels of UFM1 immunoreactivity (Figure 6H, 6I), consistent with lower levels of free UFM1 in lysates from neurons with Tau inclusions. Our findings support that UFMylation promotes Tau pathology in AD.

## DISCUSSION

Here we report a robust 4R tauopathy model in iPSC-derived human neurons, recapitulating seeding-induced propagation of Tau inclusions in human tauopathies with endogenous levels of Tau. Our model extends the recent progress in modeling 4R tauopathy in human neurons (Capano et al., 2022; Manos *et al.,* 2022) and provides a unique platform to investigate the cellular machinery underlying Tau propagation. Compared with other iPSC models with limited somatic inclusions (Manos *et al*., 2022), our pure 4R tauopathy model exhibits robust somatic 4R tau inclusions upon seeding of Tau fibrils in a time-dependent manner. Along with fibrillar Tau structures, we observed striking subcellular alterations in the soma of inclusion-bearing neurons, reminiscent of those present in AD brains. Different from the direct programming of fibroblasts by viral-mediated transduction (Capano *et al*., 2022), the isogenic iPSC lines were engineered via CRISPR editing of splicing sites to express 4R-Tau or 4R-P301S-Tau, which allows the production of large quantities of neurons homogeneously and reproducibly, facilitating the application of CRISPRi/a screenings.

Given the close association of the progression of Tau aggregation with cognitive decline, there is an urgent need to model the propagation of tauopathy in human neurons. Despite obvious differences between our *in vitro* Tau inclusion model and tangle-bearing neurons in AD brains, significant transcriptional overlaps were observed, highlighting the pathophysiological relevance of our iPSC-derived tauopathy model. Moreover, TEM analyses of inclusion-bearing neurons revealed a striking accumulation of MLBs, consistent with the endolysosomal dysfunction implicated in tauopathy and other neurodegenerative conditions. However, unlike Tau inclusions in AD, PSP, and CBD, which consist of wild-type tau, 4R neurons lacking P301S mutation exhibited no Tau inclusions in our current experimental condition. There are many possible explanations, including the challenges of modeling a chronic process that takes decades in a culture lasting only weeks and the fact that P301S mutation likely accelerates the process greatly. Indeed, transcriptomic analyses revealed that P301S mutation by itself markedly downregulates genes involved in intracellular and membrane trafficking, which could render 4R-P301S neurons more vulnerable to Tau proteostasis imbalance triggered by seeding. Consistent with this notion, functional knockdown of the v-SNARE lysosome-associated protein, VAMP7, promoted Tau propagation. Our platform enables systematic investigation on how strain selectivity, and/or selective vulnerability in cellular machinery, contributes to the heterogeneity of tauopathies.

The spread of Tau pathology often follows a predictable, trans-neuronal pattern, consistent with the role of neuronal activity in Tau spread (Franzmeier et al., 2022; Vogel et al., 2020). Our tauopathy platform enabled the tracking of Tau pathology in live human neurons and uncovered intriguing crosstalk between neuronal activity and Tau propagation. FTDP-17 patients carrying P301S mutation exhibit neuronal hyperexcitability (Garcia-Cabrero et al., 2013; Sperfeld et al., 1999). Other FTD mutations, such as V337M, also induce hyperexcitability in human neurons, suggesting a common pathogenic phenotype (Sohn *et al*., 2019). By chronically silencing neuronal activity using inhibitory DREADDs, we found that Tau propagation is significantly ameliorated, providing direct evidence that hyperexcitability associated with P301S mutation could promote Tau propagation.

Despite the correlation between the spread of Tau aggregates with cognitive decline, the exact toxic species of Tau remain poorly understood (Gendron and Petrucelli, 2009; Guerrero-Munoz et al., 2015; Spires-Jones et al., 2009). It is challenging to measure neuronal function while simultaneously monitoring their Tau pathology, as it often requires antibody staining of fixed neurons. By Halo-Tagging 4R-P301S tau, we directly compared calcium influx of live neurons with or without somatic inclusions, providing the first direct evidence that Tau inclusions impair neuronal activity in human neurons. Our observation challenges the view that Tau aggregates are not toxic *per se* and is consistent with the negative correlation of tauopathy measured by PET Tau tracer with neuronal activity measured with FDG PET (Zhang et al., 2021). This platform opens the door for a more detailed functional characterization of the effects of different Tau species in human neurons.

The cellular machinery underlying the spread of Tau aggregates remains poorly understood, partially due to a lack of an authentic robust Tau propagation model in human neurons. Our current model enables systematic analyses of cellular machinery using functional genomics. In a CRISPRi screen of a curated set of >1000 genes implicated in Tau pathobiology, we uncovered both known and unknown modifiers of Tau propagation. For example, one of the key steps of Tau propagation that remain poorly understood is how Tau seeds eventually escape the endocytic compartment and seed the aggregation of naive Tau in the cytoplasm. Our CRISPRi screen revealed that the top genes whose knockdown increases inclusions are those involved in almost all aspects of vesicular trafficking and autophagy, including those involved in the endolysosomal biogenesis and trafficking, endosome-to-lysosome fusion, Golgi trafficking, retrograde transport of vesicles from endosomes to TGN, as well as a component of retromer complex required to transport cargos from endosome to TGN or plasma membrane. These findings suggest that any deficiency in vesicular trafficking could promote Tau propagation. Based on our CRISPRi screen, we propose that the rate-limiting factor in propagation is the breaching of the organelle membranes and the escape of seeds from endocytic compartments, enabling the templating of naive Tau in the cytosol (Carosi et al., 2021).

As one of the core components of the retromer complex, VPS29 serves as a hub for vesicular trafficking (Banos-Mateos et al., 2019; Ye et al., 2020) and emerged as the top hit in this targeted screen. The two other components of the retromer complex, VPS35, whose mutation is linked with Parkinson’s disease (Williams et al., 2017), and VPS26 are not included in the library. We validated the profound modifying effects of VPS29 by establishing two clones of 4R-P301S cells lacking VPS29. VPS29 deficiency markedly enhanced seeding-induced propagation by 2–3-fold. Retromer activity is diminished in the entorhinal cortex of late-onset AD patients (Small et al., 2005), consistent with data supporting the entorhinal cortex as one of the most vulnerable regions for Tau accumulation (Llamas-Rodriguez et al., 2022). In a *Drosophila* model that overexpresses human Tau (hTau), reduction of retromer activity induces a potent enhancement of Tau toxicity, including synapse loss, axon retraction, and survival via the production of a C-terminal-truncated isoform of hTau (Asadzadeh et al., 2022). Alterations in retromer activity induced by VPS35 with D620N mutation promote Tau aggregation in the forebrain of a mouse model of Parkinson’s disease (Chen et al., 2019). In agreement with our findings, a deficiency in retromer complex activity exhibit increased levels of pathological Tau species in the cerebrospinal fluid (Simoes et al., 2020).

The proof-of-principle CRISPRi screen also uncovered novel modifiers of Tau propagation. While *MAPT* is among one of the top hits whose inhibition reduces Tau inclusions as expected, many of the top hits are mitochondrial genes, including many components in complex I, complex II, and cytochrome C. Interestingly, some of these genes overlap with the mitochondria-interacting proteins modified by FTD mutations in previous studies (Tracy *et al*., 2022). The mechanisms underlying the reduced propagation remain unknown. Since reduced neuronal activity ameliorates Tau propagation, lowering mitochondrial proteins might suppress neuronal activity, thus blocking the propagation.

Out of the top hits that promote Tau propagation are five out of the six genes in the UFMylation cascade, a process mediated by an enzyme cascade involving UBA5 (E1), UFC1 (E2), and an ER-localized trimeric ligase complex (E3) composed of UFL1, DDRGK1 (also known as UFBP1), and CDK5RAP3 (Gerakis et al., 2019). Validating the CRISPRi screen, we showed that inhibiting UFMylation by reducing UBA5 and UFM1 with shRNA markedly blocked Tau propagation. Using an antibody that preferentially binds to free UFM1, we showed that free UFM1 is reduced in inclusion-bearing neurons in our human 4R tauopathy model and tangle-bearing neurons in human AD brains. These findings provide the first evidence of involvement of UFMylation in tauopathy and that targeting UFMylation could be beneficial in reducing Tau propagation. UFMylation modulates several biological processes, and its deficiency leads to neurodevelopmental disorders (Nahorski et al., 2018). One of the principal targets of UFMylation is RPL26 (Walczak et al., 2019), whose UFMylation activates translocation-associated ribosomal quality control (Wang et al., 2023). Genetic inactivation of UFM1 or UFMylating enzymes causes the accumulation of translocation-stalled proteins at the ER and triggers ER stress. UFMylation is also strongly implicated in ER-phagy, a selective process that helps maintain ER homeostasis and cellular health by removing and recycling damaged, dysfunctional, or excessive ER components (Liang *et al*., 2020). Further studies are needed to determine if UFMylation-enhanced Tau propagation is mediated by ER stress/ER-phagy, ribosomal quality control, or other unknown mechanisms.

In summary, we developed and characterized a 4R tauopathy human iPSC platform and identified known and novel genetic modifiers. We anticipate our novel 4R-P301S iPSC lines will serve as a powerful platform to reveal the mechanisms underlying 4R tauopathies and identify new targets for drug development.

## LEAD CONTACT AND MATERIALS AVAILABILITY

The plasmids and cell lines generated in this study are available on request upon the completion of a Material Transfer Agreement (MTA). Further information and requests for resources and reagents should be directed to and will be fulfilled by the Lead Contact, Li Gan.

## EXPERIMENTAL MODEL AND SUBJECT DETAILS

### Cell cultures

To generate a stable 4R-Tau (4R) iPSC line, we used human iPSCs described in a previous study (Wang *et al*., 2017) that were engineered for inducible expression of Neurogenin2 (*NGN2*) transgene integrated into the AAVS1 locus of WTC11 cells with a wild-type genetic background (Miyaoka et al., 2014). To force the inclusion of exon 10 that allows the predominant expression of 4R tau under the regulation of the endogenous *MAPT* transcription unit, a donor plasmid containing several mutations around exon 10 was made. The mutated exon 10 region and a puromycin selection cassette were flanked by the left and right homology arms obtained from WTC11 *MAPT* genomic sequences. Two sgRNAs were selected based on the IDT CRISPR design online tool. Two guide RNAs and Cas9 protein (IDT) were incubated for 10 min at room temperature. RNP was co-electroporated with 1.8 µg of donor DNA into WTC11 iPSCs (0.3 × 10^6^) using the Lonza Nucleofector system (Lonza). Cells were then seeded in a 12-well plate. After 48 hours, transfected cells were selected with puromycin for 5 days. Knock-in clones were isolated by splitting single cells into 96-well plates. To identify the knock-in monoclonal cells, primers were designed to flank the outside of the homology arms and to be on the transgene of the targeting vector. Genomic DNA was isolated from individual targeted clones grown on 96-well plates. Homologous recombination events were identified by two simple PCR screenings. The PCR products from the positive clones were validated by Sanger sequencing. Non-specific integration and off-target were performed by PCR with the genomic DNA samples, followed by Sanger sequencing. Homozygous clones were further characterized by PCR and expanded for future use.

To excise the FRT-flanked puromycin cassette from the targeted alleles, 2 µg of pCDH-EF1-FLPe DNA was electroporated into 4R iPSCs (0.3 × 10^6^) using the Lonza Nucleofector system (Lonza). Cells were then seeded in a 12-well plate. After 48 hours, FRT cassette-deleted clones were isolated by splitting single cells into 96-well plates. To characterize the deletion driven by FLP-mediated recombination, primers flanking the FRT cassette were designed. Genomic DNA was isolated from individual clones grown on 96-well plates. After FLP recombination, the FRT-EF1-Puro-T2A-eGFP-FRT cassette was excised, leaving a single FRT that was identified by simple PCR screening. The excised locus yielded a 734-bp fragment, whereas the original locus containing the FRT cassette showed a 3.1-kb fragment. The 734-bp PCR products from the homozygous clones were validated by Sanger sequencing. Homozygous clones were further characterized by PCR and expanded for future use.

To further generate a stable disease-associated *MAPT*-P301S (4R-P301S) model, a long single-stranded DNA (ssDNA) donor template was designed to introduce C>T to obtain the intended point mutation that leads to the change of proline 301 to serine on Tau. The point mutation was flanked by 60-bp left and right homology arms that were obtained from the WTC11 genomic sequence surrounding the mutation. A sgRNA was designed at the mutated region to prevent recutting after homology-directed recombination. RNP and ssDNA (Ultramer DNA oligos, IDT) were electroporated into 4R iPSCs and selected as described above. Homologous recombination events were identified by PCR with two primers outside of the homology arms, followed by Sanger sequencing to determine the integration of the correct mutations and the absence of any additional unwanted mutations surrounding the site. Homozygous clones were further characterized by PCR and expanded for future use.

To knock out *MAPT* in the NGN2 iPSC genome, two sgRNAs were designed to delete a ∼1.8-kb fragment on the *MAPT* locus. One sgRNA specifically recognized the upstream of the *MAPT* promoter, and another sgRNA specifically recognized the exon containing the ATG start codon of the *MAPT* gene. RNP was electroporated into NGN2 iPSCs as described above. To validate the deletion, primers flanking the outside of the two sgRNAs target sequences were designed, and positive clones were screened by PCR. Homozygous clones were further characterized by PCR with three primers, two primers outside of the two sgRNAs target sequences, and one primer within the deleted region. The deletion and locus integrity were validated by Sanger sequencing of the PCR products of the homozygous clones. Homozygous *MAPT* KO clones were expanded for future use.

To generate a stable line expressing endogenous 4R-P301S-HaloTag-Tau fusion protein, a donor plasmid containing HaloTag cDNA fused with the exon containing ATG start codon of the *MAPT* gene was made. The donor construct consists of HaloTag-5′ Tau and a puromycin selection cassette that were flanked by left and right homology arms obtained from WTC11 *MAPT* genomic sequences. An sgRNA was selected using the IDT CRISPR design online tool. RNP and 1.8 µg of donor DNA were electroporated into 4R-P301S iPSCs using the Lonza Nucleofector system (Lonza) and selected as described. Integration of HaloTag-5′ Tau at the target locus was verified by two PCRs using primers flanking the outside of the homology arms and on the transgene of the donor construct. The PCR products from the positive clones were validated by Sanger sequencing. Non-specific integration was identified. Homozygous clones were further characterized by PCR and expanded for future use.

To knock in the stable CRISPRi constitutive system into 4R-P301S iPSC genome, 0.3 x 10^5^ iPSCs were electroporated with 0.5 µg of each TALEN DNA plasmid and 1 µg of pC13N-dCas9-BFP-KRAB (Addgene, 127968) donor DNA using the Lonza Nucleofector system (Lonza) and selected as described. Cells were then seeded in a 12-well plate. After 48 hours, transfected cells were expanded and seeded in 6-well plates and further expanded. iPSCs were dissociated using Accutase to prepare single-cell suspensions, followed by single-cell sorting based on the BFP expression. Sorted cells were recovered in a 12-well plate for 2 days with ROCK inhibitor (Y-27632, Cayman chemicals) and then isolated by splitting single cells into 96-well plates. To identify the dCas9 clones, primers on the *CLYBL* locus flanking the outside of the homology arms and primers on the transgene of the targeting vector were designed. Genomic DNA was isolated from individual targeted clones grown on 96-well plates. Homologous recombination events were identified by two simple PCR screenings. The PCR products from the positive clones were validated by Sanger sequencing. Non-specific integration was identified and homozygous 4R-P301S-dCas9 clones were further characterized by PCR and expanded for future use.

To stably knock out the *VPS29* gene in the 4R-P301S iPSC genome, two sgRNAs were designed to delete an ∼8-kb fragment on the *VPS29* locus. One sgRNA specifically recognized the upstream of the *VPS29* promoter, and another sgRNA specifically recognized the downstream of exon 3. RNP was electroporated into 4R-P301S iPSCs as described. To screen the deletion, primers flanking the outside of the two sgRNAs target sequences were designed and positive clones were screened by PCR. With efficient CRISPR cutting and repair of DNA through non-homologous end joining, a ∼481-bp product was expected for the deletion. Positive clones were further characterized by PCR with three primers, two primers outside of the two sgRNAs target sequences, and one primer within the deleted region. Monoallelic targeting was indicated by the appearance of two distinct products of wild-type size and targeted size, the wild-type locus yielded a 207-bp fragment, whereas the targeted locus yielded a 400-bp product. A single fragment of 400 bp indicates that both alleles were targeted. The deletion and locus integrity were validated by Sanger sequencing of the PCR products of the homozygous clones. Homozygous clones were expanded for future use. All genotyping primers are listed in Table S1.

Pre-differentiation of human iPSCs into neurons was initiated by plating 1.5 x10^6^ iPSCs in one well of Matrigel-coated 6-well plates with Knockout DMEM/F-12 medium containing doxycycline (2 µg/mL), N2 supplement, non-essential amino acids, brain-derived neurotrophic factor (10 ng/mL, Peprotech), neurotrophin-3 (10 ng/mL, PeproTech) and ROCK inhibitor (Y-27632, Cayman chemicals). The medium was replaced the next day without ROCK inhibitor, and pre-differentiation was maintained for a total of 3 days. On day 0, the pre-differentiated precursor cells were dissociated with Accutase and re-plated generally onto laminin-coated coverslips (Neuvitro) at 1.5 x10^5^ cells/coverslip or 12-well tissue culture plates (Corning) at 3.8 x10^5^ cells/well for the growth of neuronal cultures in Neurobasal Plus medium containing B27 supplement, Glutamax, BDNF (10 ng/mL) and NT3 (10 ng/mL) with doxycycline (2 µg/mL). Half of the medium was replaced on day 3, as well as on day 7 with the removal of doxycycline from the fresh medium. The medium volume was doubled on day 21. Thereafter, one-half of the medium was replaced with fresh medium weekly until the cells were collected. All human iPSC lines used in this work have been regularly tested for mycoplasma (Lonza) and karyotyped (MSK Molecular Cytogenetics Core).

### Human subjects

The tissues used for this study were the mid-frontal cortices from brains of age-matched patients with AD (n=7, 3 females and 4 males). Samples were obtained from the University of Pennsylvania brain bank. All brains were donated after consent from the next-of-kin or an individual with legal authority to grant such permission. Brain tissues of University of Pennsylvania brain bank used in this study are not considered identified “human subjects” and are not subject to IRB oversight. The institutional review board has determined that clinicopathologic studies on de-identified postmortem tissue samples are exempt from human subject research according to Exemption 45 CFR 46.104(d)(2). Additional information about the donors can be found in Table S9.

## METHOD DETAILS

### Preparation and seeding of recombinant K18-P301L Tau fibrils

Myc-tagged K18-P301L Tau was expressed and purified as described (Mok et al., 2018). Briefly, protein expression was induced in Terrific Broth containing the chemical chaperone betaine (10 mM) and IPTG (500 μM) for 3 hours at 30°C. Tau was purified via the following major steps: mechanical lysis, boiling, centrifugation, and cation exchange. Purified Tau fractions were dialyzed into aggregation assay buffer (PBS pH 7.4, 2 mM DTT). To minimize potential endotoxin contamination, purified Tau was incubated with poly(epsilon-lysine)-conjugated resin (Pierce), and then tested post-treatment for endotoxin levels using the ToxinSensorTM Chromogenic LAL Endotoxin Assay Kit (Genscript). Endotoxin levels of tau were < 0.1 EU/mL at working concentrations. Purified Tau aliquots were stored at -80°C prior to aggregation. To induce Tau aggregation, 88 µg/mL of freshly prepared heparin sodium salt (Santa Cruz Biotechnology) was added to K18-P301L Tau (20 μM) in aggregation assay buffer. Aggregation was carried out in low-retention 1.7 mL microcentrifuge tubes at 37 °C with shaking at 800 rpm for 24 hours. Aggregated Tau was isolated by centrifugation at 100,000xg for 1 hour at 4 °C. Pelleted Tau aggregates were resuspended in PBS (pH 7.4) with sterile plastic pestles and stored in low-binding tubes (CoStar) at -80 °C. Tau fibril preparations were retested to confirm endotoxin levels < 0.1 EU/mL at working concentrations. The concentration of aggregated Tau was quantified by Pierce BCA assay. Fibrils were thawed on ice and mixed by pipetting, and the volume needed for seeding was transferred to 100 μL of sterile DPBS. The fibrils were sonicated at 4°C, 10 minutes on/off, 30-second pulse, and amplitude of 40% using a water bath sonicator (EpiSonic 2000, EpigenTek). The fibrils were added to appropriate media volume in wells with neurons to achieve a 1.5 or 3 µg/mL final concentration.

### Western blot

Human iPSC-derived neurons were washed twice with cold DPBS, centrifuged at 300g for 5 minutes at 4°C and homogenized in cold N-PER Neuronal Protein Extraction Reagent (Thermo Fisher) or cold RIPA lysis buffer (Thermo Fisher) supplemented with protease inhibitor cocktail (Millipore Sigma), phosphatase inhibitor cocktail (MilliporeSigma) and deacetylase inhibitors, including nicotinamide (Millipore Sigma) and trichostatin A (Millipore Sigma). N-PER and RIPA lysates were incubated on ice for 10 minutes, centrifuged at 14,000 g for 10 minutes at 4°C, and the supernatants were collected, according to the manufacturer’s instructions. For Triton X-100 soluble/insoluble fractionation, cell pellets were resuspended in cold lysis buffer containing 50 mM Tris pH 7.6, 150 mM NaCl, 1% (v/v) Triton-X 100, protease inhibitor cocktail (Roche), phosphatase inhibitor cocktail (Sigma), and deacetylase inhibitors, including nicotinamide and trichostatin A. Samples in lysis buffer were incubated on ice for 30 minutes, centrifuged at 20,000 g for 30 minutes at 4°C, and the supernatant was collected as a Triton-soluble fraction. Triton-insoluble pellets were further resuspended in lysis buffer containing 50 mM Tris, pH 7.6, 150 mM NaCl, 5% (w/v) sodium dodecyl sulfate (SDS), protease inhibitor cocktail (Roche), phosphatase inhibitor cocktail (Sigma), nicotinamide and trichostatin A using one-third of the final volume of the Triton-soluble fraction. Insoluble samples were sonicated at 16°C for 4 minutes on/off with 2-second pulses and 20% amplitude using a water bath sonicator (EpiSonic 2000, EpigenTek), centrifuged at 20,000 g for 30 minutes at 20°C and the supernatant was collected as Triton-insoluble fraction. Protein concentration was determined using a Pierce BCA Protein Assay Kit (Thermo Fisher) for all samples except the Triton-insoluble fractions. Equal amounts of protein for each sample were run on a 4–12% SDS-PAGE gel (Invitrogen). Nitrocellulose membranes (GE Healthcare) were used for the transfer of proteins, and blots were blocked with 5% milk in TBS-Tween. Primary and secondary antibodies were diluted and incubated in the blocking solution. Primary antibodies used included mouse anti-tau (HT7, Thermo Fisher), anti-Tau 3-repeat isoform (RD3, Sigma), anti-Tau 4-repeat isoform (ET3, Sigma), anti-phosphorylated Tau (AT8, Thermo Fisher), anti-phosphorylated PHF Tau (AT180, Thermo Fisher), anti-phosphorylated PHF Tau (AT270, Thermo Fisher), and rabbit anti-UFM1 (Abcam), anti-GAPDH (GeneTex). Secondary HRP antibodies (Millipore) and chemiluminescence (BioRad) were used for detection of immunoblotting and bands in immunoblots were quantified by intensity using ImageLab (BioRad) or FIJI (NIH) software.

### Immunocytochemistry and imaging

For human iPSCs and iPSC-derived neurons, cells were fixed in 4% paraformaldehyde (Electron Microscopy Sciences) diluted in PBS for 15 minutes and washed three times for 5 minutes in DPBS with Mg^2+^ and Ca^2+^. Cells were permeabilized in 0.1% Triton X-100 diluted in DPBS for 10 minutes and then placed in a blocking solution containing 0.1% Triton X-100 and 5% goat serum in DPBS for 1 hour at room temperature. Primary antibodies diluted in blocking solution were added to the cells overnight at 4°C and then washed three times for 5 minutes with DPBS. Primary antibodies used included: chicken anti-MAP2 (Novus Biologicals), mouse anti-conformationally abnormal Tau (MC1, Peter Davies), anti-phosphorylated Tau (AT8, Thermo Fisher), anti-SSEA (Abcam), anti-TRA-1-60 (Abcam), anti-TRA-1-81 (Abcam), and rabbit anti-oligomeric Tau (TTC18, Rakez Kayed), anti-UFM1 (Abcam), anti-GFP (Abcam), anti-OCT4 (Abcam), NANOG (Abcam), and SOX2 (Cell Signaling). Secondary antibodies diluted in blocking solution included donkey anti-mouse, anti-rabbit, anti-chicken and anti-human antibodies conjugated with Alexa Fluor 488, Alexa Fluor 568, or Alexa Fluor 646 fluorophores (1:500, Life Technologies), and they were added to the cells for 1 hour at room temperature. The coverslips were washed three more times for 5 min with DPBS before being mounted on slides with Vectashield containing DAPI (Thermo Fisher). Images were acquired using Apotome (Zeiss), Keyence (BZ-X710), or LSM 880 Laser Scanning Confocal Microscope (Zeiss) and processed with Zen 3.2 software. The settings used for image acquisition were selected to keep the majority of the brightest pixel intensities from reaching saturation. Quantification was performed using FIJI (NIH) software.

For human AD brain tissue, sections were formalin-fixed and paraffin-embedded into glass slides. Slides were placed for 10 minutes at 60 °C in an oven and then de-paraffinized by washing with xylene three times for 5 mins, 100% ethanol two times for 2 mins, 95% Ethanol for 2-5 mins, and deionized water 3 times for 2 minutes. For antigen retrieval, the slides were placed in working Tris-EDTA buffer (10 mM Tris base, 1 mM EDTA solution, 0.05% Tween 20, pH 9.0) and placed in a pressure cooker (Cuisinart) at high pressure for 15 minutes. The slides were cooled to RT and washed with cool deionized water three times for 2 minutes. Sections were washed with 1x PBS three times for 2 minutes and incubated with 1x TrueBlack in 70% Ethanol for 30 seconds. The reaction was stopped by placing the slides in 1x PBS and further washing with 1x PBS three times for 2 minutes. Sections were blocked with 5% donkey serum in 1x PBS for 1 hour and incubated with UFM1 (Abcam 1:100) and MC1 (1:1000) diluted in 1% donkey serum in 1x PBS solution overnight in a humidified slide chamber. The following day, the slides were washed in 1x PBS three times for 2 minutes and incubated with AF 488 donkey anti-mouse (1:500) and AF 568 donkey anti-rabbit (1:500) diluted in 1% donkey serum in 1x PBS solution for 1 hour in a humidified slide chamber. The slides were washed in 1x PBS three times for 2 minutes and incubated with Hoescht 33342 diluted (1:1000) in 1x PBS for 15 minutes. The slides were washed in 1x PBS three times for 2 minutes and covered with Vectashield mounting medium without DAPI (Vector Labs). Five imaging fields in the gray matter of the same case were captured using a fluorescent microscope (Keyence BZ-X710), and UFM1 mean intensity of n=3 MC1+ and MC1-cells was quantified using NIS-Element Analysis software.

### Flow cytometry

For flow cytometry, briefly, human iPSC-differentiated neurons were dissociated from the plate with papain solution (20 U/mL papain and 5 mM MgCl_2_ in HBSS) at 37 °C for 30 mins. Papain was quenched with 3x volume DMEM with 10% FBS. Cells were fixed with zinc fixation buffer (0.1 M Tris-HCl with pH 6.5, 0.5% ZnCl_2_, 0.5% Zn acetate and 0.05% CaCl_2_) overnight at 4 °C. The next day, samples were washed twice with TBS and resuspended in permeabilization buffer (10% donkey serum, 10% 10X TBS, 3% BSA, 1% glycine, 0.5% Tween-20) for 15 minutes. Primary antibodies are added into permeabilization buffer by pipetting up and down to separate cells into a single-cell suspension, and samples were incubated either at 4°C overnight or at room temperature for 1 hour. The samples were incubated in permeabilization buffer with secondary antibodies at room temperature for 1 hour after washing twice in TBS. Samples were washed twice with TBS, analyzed with Attune NxT Flow Cytometer (Thermo Fisher) or LSRFortessa cell analyzer (BD), and the data were processed using FlowJo v10 software. To detect endogenous Tau inclusions, anti-MC1 (1:200, Peter Davies) primary antibody was used.

### Bulk RNA-sequencing

4R and 4R-P301S iPSCs were differentiated into neurons on 12-well tissue culture plates, and half of the samples were seeded with 1.5 µg/ml K18 on D8 as described. Neurons were maintained and collected on D43. Total RNA was extracted from the samples using QuickRNA MicroPrep Kit (Zymo Research). After RNA isolation, total RNA integrity was checked using a 2100 Bioanalyzer (Agilent Technologies), and concentrations were measured by Nanodrop (Thermo Fisher). Preparation of the RNA sample library and RNA-seq were performed by the Genomics Core Laboratory at Weill Cornell Medicine using Illumina Stranded mRNA Sample Library Preparation kit (Illumina), according to the manufacturer’s protocol. The normalized cDNA libraries were pooled and sequenced on NovaSeq 6000 (Illumina) with pair-end 100 cycles. The raw sequencing reads in BCL format were processed through bcl2fastq 2.19 (Illumina) for FASTQ conversion and demultiplexing. Paired-end reads were cleaned by trimming adapter sequences and low-quality bases using cutadapt v1.18 (Martin, 2011) and aligned and mapped to the human genome (GRCh38) using STAR v2.5.2 (Dobin et al., 2013). The transcriptome reconstruction was performed by Cufflinks v2.1.2, and the abundance of transcripts was measured with Cufflinks in Fragments Per Kilobase of exon model per Million mapped reads (FPKM) (Trapnell et al., 2010). Raw read counts per gene were extracted using HTSeq-count v0.11.2 (Anders et al., 2015). DEGs were identified using the RStudio DESeq2 package (Love *et al*., 2014), and plots were generated with the ggplot2 package v3.4.1. To find enriched annotations within hit genes, Gene Ontology analysis was performed using Gene Set Enrichment Analysis software on the four conditions using the Molecular Signatures Database C5 ontology gene sets (Liberzon et al., 2011; Subramanian et al., 2005).

### Single-cell RNA-sequencing

4R-P301S iPSCs were differentiated into neurons, K18-seeded, and matured to D39 as described. Neuronal dissociation was performed based on established protocols (Jerber et al., 2021; Tian *et al*., 2019). Neurons were washed twice with 1X DPBS (Thermo Fisher) before adding a 1:1 Accutase (Thermo Fisher Scientific) and 1X DPBS solution containing 20 U/ml papain (Worthington Biochemical) and 50 μg/mL DNase I (Worthington Biochemical). The cells were incubated at 37 °C for up to 35 minutes before adding four times the dissociation solution volume of quenching solution composed of DMEM/F12 (Thermo Fisher Scientific), 10% FBS (Thermo Fisher Scientific), 10 µM ROCK inhibitor (Y-27632, Cayman chemicals) and 50 μg/mL DNase I (Worthington Biochemical) and transferring the neurons coming off as a layer to a 15-mL tube. Cells were centrifuged at 200x g for 4 minutes at RT, resuspended in quenching solution, gently dissociated using a P1000, and collected in a 15-mL tube capped with a 40-µm cell strainer (Corning), and washed three additional times in 1X DPBS containing 0.04% BSA (Sigma Aldrich). Single-cell suspensions were counted using an automated cell counter (TC20, BioRad). 10x Genomics 3’ library construction was prepared with Chromium Single Cell 3’ Reagent Kits v3.1 (10x Genomics, PN-1000268) according to the manufacturer’s protocol. cDNA and library fragment analysis were performed using the Agilent Fragment Analyzer systems. The libraries were sequenced on the NovaSeq 6000 sequencer (Illumina) with PE 2 x 50 paired-end kits by using the following read length: 28 cycles Read 1, 10 cycles i7 index, 10 cycles i5 index, and 90 cycles Read 2.

### Transmission electron microscopy

Neurons were washed once in DPBS and fixed overnight at 4°C with 2.5% glutaraldehyde and 4% PFA in 0.2 M sodium cacodylate buffer (pH 7.3) supplemented with 5% saturated aqueous solution of picric acid (Ito and Karnovsky, 1968). The next day, cells were washed three times for 10 minutes each with 0.1 M sodium cacodylate buffer (pH 7.3), postfixed in 1% osmium tetroxide and 1.5% potassium ferricyanide for 1 hour at room temperature, dehydrated in graded alcohols (50, 70, 85, 95%, and three times 100%) for 15 minutes at each concentration, and embedded in epoxy resin. Thin sections (65 nm) were cut on 200-mesh copper grids, stained with 1.5% uranyl acetate, followed by lead citrate (Venable and Coggeshall, 1965), and examined using a digital TEM (JSM1400, JEOL Ltd). Images were acquired using a Veleta 2k×2k charge-coupled device camera (Olympus-SIS) at low magnifications to capture the soma or dendrites and then at 150,000x to capture tau fibrils. Multilamellar bodies (MLBs) were identified using morphological criteria. Both the number of MLBs per soma and the total area of MLBs normalized to the soma area were calculated by manual contouring in Fiji (NIH).

### Calcium imaging

4R-P301S-HaloTag iPSCs were differentiated into neurons on coverslips as described. Neurons were transduced with hSyn-jGCaMP8f lentivirus on D6. This lentiviral construct was made using the 3^rd^ generation lentiviral plasmid FUGW (Addgene, 14883), where the PacI + EcoRI fragment was replaced by the hSyn-GCaMP8f fragment (Zhang *et al*., 2023). pGP-AAV-hSyn-jGCaMP8f-WPRE was a gift from GENIE Project (Addgene, 162376). The medium was changed on D7, and neurons were seeded with 1.5 µg/mL K18-tau fibrils. Neurons were maintained as described until imaging at 4 and 5 weeks old. On the day of imaging, JFX-549 HaloTag ligand (HHMI Janelia) was diluted into fresh maturation medium to 1 μM, and one-fifth of neuronal medium was replaced with diluted ligand (200 nM final ligand concentration). Cells were incubated at 37 °C for 15 minutes. An equal volume of medium was replaced with half-fresh medium, and half-conditioned medium from a control well. At the time of imaging, the coverslip was gently washed and placed into a glass-bottom chamber (RC-26G, Warner Instruments) containing Ca^2+^ imaging buffer (20 mM HEPES, 119 mM NaCl, 5 mM KCl, 2 mM MgCl_2_, 30 mM glucose, 2 mM CaCl_2_, pH 7.2–7.4).

The temperature was maintained at 37 °C by a dual chamber heat controller (TC-344C, Warner Instruments). Fluorescence time-lapse images were collected on a Nikon FN1 microscope using a 60x, 1.0 NA objective (CFI APO 60XW NIR, Nikon) and a C-FL GFP filter cube. An X-CITE LED illuminator (Nikon) was used for excitation. Images were collected using an ORCA-Fusion CMOS camera (Hamamatsu) with 4×4 binning (576×576 pixel resolution, 16-bit grayscale depth, 0.43 μm/pix) and NIS-Elements AR software (Nikon). HaloTag signal per field of view was collected using a C-FL-DS RED filter cube (Nikon). Exposure time was set to 20 milliseconds. For spontaneous activity, 4–5 fields per coverslip were acquired at 30 Hz for 2 minutes. After imaging spontaneous activity, one field per coverslip was imaged for 10 minutes at 20 Hz to minimize photo-bleaching immediately after 50 mM KCl perfusion.

### DREADDs neuronal silencing

4R-P301S-HaloTag iPSCs were differentiated into neurons on coverslips as described. Neurons were transduced with hSyn-hM4Di-mCherry lentivirus on D6. This lentiviral construct was made using the 3^rd^ generation lentiviral plasmid FUGW (Addgene, 14883), where the PacI + EcoRI fragment was replaced by hSyn-hM4D(Gi)-mCherry. pAAV-hSyn-hM4D(Gi)-mCherry was a gift from Bryan Roth (Addgene, 50475; http://n2t.net/addgene). The medium was changed on D7 and neurons were seeded with 1.5 µg/mL K18-tau fibrils. Neurons were maintained as described. Beginning at D14, clozapine-N-oxide (CNO; Tocris) was included in weekly media changes for a final concentration of 10 μM. Neurons were fixed and stained with MC1 on D29. For MC1 quantification, stained neurons were imaged with a fluorescence microscope (Keyence BZ-X710). A 20X objective was used to acquire a series of 10 Z-stack images per coverslip. To validate the neuronal silencing effect of CNO by calcium imaging, 4R-P301S neurons were differentiated and transduced with hSyn-hM4Di-mCherry and hSyn-GCaMP8f lentivirus on D6. At D36, JFX-549 was added to neurons and prepared for imaging as described above. Exposure time was set to 20 milliseconds. Neurons were perfused with 10 μM CNO at 60 seconds and calcium traces in one field were recorded for 400 seconds.

### VAMP7^DN^ human Tau ELISA and immunocytochemistry

For Human Tau ELISA, 4R and 4R-P301S iPSCs were differentiated on 12-well tissue culture plates as described previously. Neurons were seeded with 1.5 µg/mL K18 on D14. Two weeks after seeding on D28, the cell culture medium was removed and replaced with high KCl extracellular solution containing (in mM): 68.5 NaCl, 50 KCl, 10 HEPES, 20 glucose, 1 MgCl_2_, 2.5 CaCl_2_ at pH 7.4. The neurons were incubated with KCl extracellular solution for 30 minutes at 37 °C. The extracellular solution was collected for ELISA analysis. The neurons were lysed with cold RIPA buffer (100 μL/well) and homogenized on ice with a handheld homogenizer for 2 minutes for each sample. Samples were centrifuged at 20,000 g for 10 minutes at 4 °C and the supernatants were collected. Protein concentrations for the extracellular solution and cell lysate supernatant were determined by BCA assay with Pierce BCA protein assay kit (Thermo Fisher Scientific) according to manufacture protocol on Synergy H1 microplate reader (Biotek). A concentration series of seven Tau-352 protein standards were prepared for human Tau antibody-coated plates with 96-wells (Human Tau (Total) ELISA Kit, Cat. No. KHB0042). The extracellular solution was diluted 1:100, and the lysate was diluted to 1:1000 for all samples. Each standard (100 μL) or sample (50 μL) plus Standard Diluent Buffer (50 μL) was added to individual blocked wells, tapped to mix, and covered and incubated for 2 hours at room temperature shaking at 100 rpm. The solution was thoroughly aspirated, and wells were washed 4x with 400 μL of 1x wash buffer. Then 100 μL human Tau (Total) biotin conjugate solution was added into each well except the chromogen blanks. The plate was covered and incubated for 1 hour at RT shaking at 100 rpm. The solution was thoroughly aspirated, and wells were washed as described above. Next, 100 μL of 1x streptavidin-HRP solution was added into each well except for the chromogen blanks. The plate was covered and incubated for 30 minutes at RT shaking at 100 rpm. The solution was thoroughly aspirated, and wells were washed as described above. Next, 100 μL of stabilized chromogen was added to each well, following 50 μL streptavidin-HRP solution (1:8000). The plate was incubated for 30 minutes at RT in the dark. Lastly, 100 μL of stop solution was added to each well, tapped to mix, and the plate was read at 450 nm. The amount of secreted Tau was quantified from the percent of extracellular Tau detected out of the total amount of intracellular and extracellular Tau for each well of culture.

For immunocytochemistry, 4R and 4R-P301S iPSCs were differentiated into neurons on coverslips as described. Neurons were transduced with GFP or GFP-VAMP7^DN^ lentivirus on D13. pFUGW-eGFP and pFUGW-eGFP-VAMP7^2-120^ were gifts from Manu Sharma (Xie *et al*., 2022) The medium was changed on D14, and neurons were seeded with 1.5 µg/mL of K18-Tau fibrils. Neurons were maintained as described, and fixed and stained with MC1 on D28. For MC1 quantification, stained neurons were imaged with a fluorescence microscope (Keyence BZ-X710). A 20X objective was used to acquire a series of 10 Z-stack images per coverslip.

### Lentivirus generation

Generally, lentiviral particles were generated for pFUGW-hSyn-GCaMP8f, pFUGW-hM4Di-mCherry, pFUGW-eGFP, and pFUGW-eGFP-VAMP7^DN^ as follows. HEK293T cells (ATCC) were thawed, passed, and expanded in T75 flasks with 15 mL of DMEM/F-12 (basal medium supplemented with 10% FBS and 1% penicillin/streptomycin). Cells were passed into 10-cm dishes (1:10), and 2 days later upon reaching ∼80% confluency, transfection mix for each plasmid was prepared in the following manner: 1 µg of transfer plasmid and 2 µg of second-generation packaging DNA mix containing psPAX (Addgene #12260) and pMD2.G (Addgene #12259) (molarity 1:1) were diluted into Opti-MEM I Reduced Serum Medium (GIBCO; Cat. No. 31985070); TransIT-LT1 (Mirus Bio, Cat. No. MIR6600) was diluted into Opti-MEM and incubated at room temperature for 5 minutes; the diluted DNA solution was added to the diluted Lipofectamine solution, inverted several times to mix, and incubated at room temperature for 10 minutes. Volumes depended on the number of 10-cm plates used according to the manufacturer’s protocol. After incubation, the transfection solution was gently added dropwise to each 10-cm dish with HEK293T cells, and the plates were briefly and gently swirled to mix. At 48 hours, HEK293T medium was transferred into 50-mL conical tubes, and the plates were replaced with fresh medium. At 72 hours, HEK293T medium was collected again and combined with previously collected media. The supernatant was carefully transferred to a syringe fitted with a 0.45-μm PVDF filter to filter the virus-containing solution into a new 50-mL conical. Cold Lenti-X Concentrator (Takara; Cat. No. 631232) corresponding to one-third volume of the supernatant was added to the filtered solution, which was then mixed well and stored at 4C for 24 hours. After incubation, the solution was centrifuged at 4 °C for 45 minutes at 1,500 g, and the supernatant was decanted. The virus-containing pellet was resuspended in 200 μL of DPBS (ThermoFisher) and stored at -80 °C. Lentiviral particles for single sgRNAs (non-targeting, TFRC, VPS29, LAMTOR5, UFM1) and shRNAs (UFM1, UBA5) were generated as described above with the following modifications: third-generation packaging DNA mix containing pRSV-REV (Addgene #12253), pMDLg/pRRE (Addgene #12251), and pMD2.G (Addgene #12259) (mixed 1:1:1); 1:4 Lentivirus Precipitation Solution (Alstem; Cat. No. VC125) for concentration step; lentiviral supernatant collection at 48 hours.

### Functional validation of 4R-P301S-dCas9 iPSC line

4R-P301S-dCas9 iPSCs were seeded at 5 x 10^5^ per well in six-well tissue culture plates with ROCK inhibitor. The following day at 24 hours, the medium was replaced with fresh medium, and iPSCs were transduced with single sgRNA lentiviruses (NTC, TFRC). The next day, we performed a complete media change. The MOI, quantified as the fraction of BFP-positive cells by flow cytometry, was ∼12%. The following day, we began 0.8 µg/mL puromycin selection (ThermoFisher) to enrich sgRNA-expressing cells. On day 2 of selection, the cells were split 1:3 into six-well plates. After 2 more days of selection, the cells were assessed by flow cytometry (∼75% expressed high levels of BFP). The following day, the iPSCs were seeded for pre-differentiation and were further differentiated, seeded with 1.5 µg/mL of K18 on D7, and maintained as described. On D21, neurons were dissociated, resuspended, and blocked for 15 minutes with 1:20 human FC block (BD Biosciences; Cat. No. 564220) and then stained with 1:66 PE-Cy7 anti-human CD71 (TFRC) (BioLegend; Cat. No. 334112) for 30 minutes in the dark. Cells were washed with DPBS before analyzing them by flow cytometry using the LSRFortessa cell analyzer (BD). Flow cytometry data were analyzed using FlowJo (FlowJo, v.10.7.1); percent BFP+/CD71+ TFRC cells were normalized to those in the NTC samples; and data were plotted as fold-change using Prism 8 (GraphPad, v.9.4.1).

### Primary CRISPRi screen

The CRISPRi custom library targeting hits identified in a genome-wide modifier screen of tau oligomer levels in human iPSC-derived neurons (Samelson *et al*., 2023) with the top five sgRNAs per gene based on the v2 CRISPRi sgRNA design algorithm (Horlbeck et al., 2016) was packaged into lentivirus for transduction of iPSCs as follows. One 15-cm dish was seeded with 12 x10^6^ HEK293T cells in 30 mL of DMEM/F-12 (basal medium supplemented with 10% FBS and 1% penicillin/streptomycin). The next day, the custom library transfection mix was prepared in the following manner: 15 µg of custom library plasmid and 15 µg of third-generation packaging DNA mix containing pRSV-REV (Addgene #12253), pMDLg/pRRE (Addgene #12251), and pMD2.G (Addgene #12259) (mixed 1:1:1) were diluted into 3 mL of Opti-MEM I Reduced Serum Medium (GIBCO; Cat. No. 31985070); 90 μL of TransIT-LT1 (Mirus Bio, Cat. No. MIR6600) was diluted into 3 mL of Opti-MEM and incubated at RT for 5 minutes; the diluted DNA solution was added to the diluted Lipofectamine solution, inverted several times to mix, and incubated at room temperature for 10 minutes. After incubation, half of the transfection solution was gently added dropwise to each 15-cm dish with HEK293T cells, and the plates were briefly and gently swirled to mix. Two days later, HEK293T media (approximately 30 mL) was transferred into a 50-mL conical tube. The supernatant was carefully transferred to a syringe fitted with a 0.45 μm PVDF filter to filter the virus-containing solution into a new 50 mL conical. Approximately 7.5 mL of cold Lentivirus Precipitation Solution (Alstem; Cat. No. VC125) was added to the filtered solution, which was then mixed well and stored at 4C for 24 hours. Following incubation, the solution was centrifuged at 4C for 30 min at 1,500 g, and the supernatant was decanted. The virus-containing pellet was resuspended in 1 mL mTeSR Plus medium and stored at -80C.

For infection with the tau custom library, one T175 Matrigel-coated flask was seeded with 5 x10^6^ 4R-P301S-dCas9 iPSCs in mTeSR Plus medium with ROCK inhibitor. The following day at 24 hours, the medium was replaced with 35 mL fresh medium plus 500 uL of the virus-containing medium was added to the cells. The next day, we performed a complete media change, adding 30 mL mTeSR Plus medium. Two days later, we began 0.5 µg/ml Puromycin selection (ThermoFisher) in the 30 mL medium, which was the medium volume and formulation used for puromycin treatment to enrich sgRNA-expressing cells. The following day, we dissociated the cells and seeded one T175 Matrigel-coated flask with 5 x10^6^ cells in 30 mL mTeSR with ROCK inhibitor. The MOI, quantified as the fraction of BFP-positive cells by flow cytometry, was ∼30%, corresponding to a library representation of ∼250 cells per library element. Puromycin treatment was increased to 0.8 µg/ml for the following three days. At the end of treatment, cells were assessed by blue fluorescence/bright field microscopy (∼75% expressed high levels of BFP) and seeded for the MC1 screen as described below.

For the MC1 screen, 3 T175 Matrigel-coated flasks were each seeded with 12 x10^6^ cells in 30 mL N2 Pre-Differentiation Medium and differentiated as previously described into 6 15-cm PDL-coated dishes (Corning), seeded 15 x10^6^ precursor cells each. On D7, the cells were seeded with 3 µg/mL K18. On D19, cells from 5 15-cm dishes (cells in 1 dish died) were dissociated by papain, fixed, and stained with MC1 as described previously. Cells were FACS-sorted into 1 mL 30% BSA (Sigma) solution using a FACSAria II (BD) based on MC1, approximately ∼2 million MC1+ cells and ∼6 million MC1-cells, corresponding to a library representation of ∼1,000 cells per library element in MC1-group and ∼333 cells per library element in MC1+ group. The cells were pelleted at 200 g for 20 minutes, the supernatant was carefully removed and stored at -20C. Genomic DNA was extracted with the NucleoSpin Blood L kit (Macherey Nagel; Cat. No. 740954.20). The sgRNA-encoding regions were amplified, pooled, and sequenced on a MiSeq sequencer (Illumina) with single read 65 cycles including 20% Phi X Sequencing Control DNA at the Genomics Core Laboratory at Weill Cornell Medicine. The raw sequencing reads in BCL format were processed through bcl2fastq 2.20 (Illumina) for FASTQ conversion and demultiplexing, based on previously described protocols (Gilbert et al., 2014; Kampmann et al., 2014; Tian *et al*., 2019).

### Primary screen validation

4R-P301S-dCas9 iPSCs were seeded at 5 x 10^5^ per well in six-well tissue-culture plates with ROCK inhibitor. The following day at 24 hours, the medium was replaced with 2 mL of fresh medium, and iPSCs were transduced with single sgRNA lentiviruses (NTC, VPS29, LAMTOR5, UFM1). The next day, we performed a complete media change. The MOI, quantified as the fraction of BFP-positive cells by flow cytometry, was ∼22%. The following day, we began 0.8 µg/mL puromycin selection (ThermoFisher) to enrich sgRNA-expressing cells. On day 2 of selection, the cells were split 1:3 into six-well plates. After 2 more days of selection, the cells were assessed by flow cytometry (∼74% expressed high levels of BFP). The following day, the iPSCs were seeded for pre-differentiation and were further differentiated, seeded with 1.5 µg/mL of K18 on D7, and maintained as described. On D21, neurons were zinc-fixed and stained with antibodies for flow cytometry as described. The samples were analyzed LSRFortessa cell analyzer (BD), and the data were processed using FlowJo v10 software. The following primary antibody was used: anti-MC1 (1:200). NucRed Live 647 ReadyProbes Reagent was used to detect and gate on intact cells.

### VPS29^-/-^ and UFMylation cascade phenotype validation

For VPS29^-/-^ validation, 4R-P301S and 4R-P301S;*VPS29*^-/-^ iPSCs were seeded at the density of 1.2 x 10^7^ per plate in 10-cm plates in pre-differentiation medium. On day 0, pre-differentiated neurons were replated at the density of 4 x 10^5^ per well in poly-D-lysine (PDL) pre-coated 12-well plates (Corning) or at the density of 2 x 10^5^ per well on laminin pre-coated coverslips (Neuvitro) in PDL pre-coated 24-well plates (Corning) in maturation medium. On day 7, iPSCs differentiated neurons were treated with 1.5 µg/mL of K18 fibrils for 2 weeks. On day 21, neurons in 12-well plates were collected for flow cytometry analysis, and neurons on coverslips were fixed for immunocytochemistry. For UFMylation cascade validation, 4R-P301S-iPSCs were seeded and differentiated as described above. On day 3, lentivirus containing control shRNA and shRNAs for UFM1 and UBA5 were added to the culture medium. The virus-containing medium was replaced with fresh medium after 24 hours. On day 7, iPSCs differentiated neurons were treated with 5 µg/mL of K18 fibrils for 2 weeks. On day 21, neurons in 12-well plates were collected for qPCR analysis and flow cytometry analysis, and neurons on coverslips were fixed for immunocytochemistry. shRNA sequences are listed in Table S7.

For flow cytometry, the cells were cultured, dissociated, and analyzed as described. The following primary antibodies were used: anti-MC1 (1:200, Peter Davies) and anti-GFP (1:400, Abcam). For immunocytochemistry, neurons were fixed and stained as described and imaged with LSM 800 confocal microscope (Zeiss). Primary antibodies were used: anti-MC1 (1:1000 Peter Davies), anti-UFM1 (1:250, Abcam), and anti-GFP (1:2000, Abcam). For qPCR, RNA was extracted from neurons using Quick-RNA™ Microprep Plus Kit (Zymo Research), according to the manufacturer’s protocol. cDNA was synthesized using the iScript Reverse Transcription Supermix for RT-qPCR (Bio-Rad) and real-time PCR was performed using SsoAdvanced Universal SYBR® Green Supermix (Bio-Rad) with primers listed in Table S8. The thermal cycling conditions were 95 °C for 10 minutes, 45 cycles of 95 °C for 15 seconds, 60 °C for 30 seconds, and 72 °C for 30 seconds. Data were collected using the BCFX384 Touch Real-Time PCR Detection System (Bio-Rad). Each sample was tested in replicate. Gene expression fold changes were calculated by the ΔΔCT method.

## QUANTIFICATION AND STATISTICAL ANALYSIS

For statistical analysis, we used GraphPad Prism 9.2.0 software or R Version 4.2.2 with packages. Data are shown as mean ± SEM. For two sample comparison, an unpaired two-tailed Student’s t-test was used to quantify the data. For three sample comparison, we used the one-way ANOVA, followed by a Dunnett post hoc test with multiple testing correction and to set up a control. For comparison of resistant and non-resistant cell lines, a one-way ANOVA and a Tukey test with multiple testing correction. For two phenotypes and two colonies’ comparison, we used two-way ANOVA analysis, followed by Šidák’s multiple comparison’s test. The statistics used in ClueGO analyses revealed enriched networks with p-values < 0.05 from right-sided hypergeometric testing with Bonferroni correction. n=2–3 independent biological replicates were used for all experiments. n.s. denotes a non-significant difference. \**p*<0.05, \*\**p*<0.01, \*\*\**p*<0.001, \*\*\*\**p*<0.0001.

### Gene overlap analysis AD brain single-cell RNA-sequencing

Differential gene expression analysis was performed using RNA-sequencing of AT8- and AT8+ (NFT-bearing) excitatory neurons from a published human AD brain dataset (Otero-Garcia *et al*., 2022) and the FindMarkers function and MAST in Seurat. Differential gene expression overlap analysis was performed using the 4R-P301S+K18 vs 4R+K18 DEGs from the bulk RNA-seq analysis, 4R-P301S+K18 vs 4R-P301S DEGs from the pseudo bulk scRNA-seq analysis, and AT8+ vs AT8-DEGs for the AD brain analysis. Gene overlap statistical significance was assessed using the GeneOverlap R package (Shen, 2022). Gene Ontology (GO) term enrichment analyses were performed using ClueGO version 2.5.9 in Cytoscape version 3.8.2 (Bindea et al., 2009; Shannon et al., 2003). Enrichment and network analyses were applied from three ontologies including GO Molecular Function, GO Biological Process, and GO Cellular Components.

### Calcium imaging analysis

For every recorded time-lapse image, we selected ROIs covering all identifiable cell bodies using a semi-automated algorithm in NIS-Elements AR (Nikon). Neurons were categorized based on HaloTag signal punctate structures corresponding to large Tau inclusions (-puncta=without inclusion, +puncta=with inclusion). Further quantifications were performed using custom-written MATLAB (Mathworks) scripts. For single-cell activity, the fluorescence time course was measured by averaging all pixels within individual ROIs. Then we use the CaPTure toolbox to extract spike activity by a rolling average method called ‘DFF’ (Jia et al., 2011; Tippani et al., 2022). A percentile-based threshold (mean + 2 standard deviations of fluorescence) was employed to detect peak events. The amplitude and frequency of each ROI was analyzed as the single-neuron activity. To determine the network activity, the synchronous firing rate of the entire cell population in the FOV was measured (Sun and Südhof, 2021). For KCl stimulation experiments, each image frame was divided by an average of all frames acquired during the first 0.5-sec window, which functioned as a baseline. Photobleaching was corrected by fitting exponentially weighted moving averages (Jia *et al*., 2011)

### Primary screen analysis

The primary screen was analyzed using the published MAGeCK-iNC bioinformatics pipeline (Tian *et al*., 2019). Briefly, raw sequencing reads from next-generation sequencing were cropped and aligned to the reference using Bowtie v.0.12.9 to determine sgRNA counts in each sample. The quality of each screen was assessed by plotting the log_10_(counts) per sgRNA on a rank-order plot. Raw phenotype scores and significance p-values were calculated for target genes, as well as for ‘negative-control-quasi-genes’ that were generated by random sampling with replacement of five non-targeting control (NTC) sgRNAs from all NTC sgRNAs. The final phenotype score for each gene was calculated by subtracting the raw phenotype score by the median raw phenotype score of ‘negative-control-quasi-genes’ and then dividing by the standard deviation of raw phenotype scores of ‘negative-control-quasi-genes’. The hit strength, defined as the product of knockdown phenotype score and –log_10_(p-value), was then calculated for all genes in the library and for ‘negative-control-quasi-genes’ generated above. Hit genes were determined based on the hit strength cutoff corresponding to a false-discovery rate (FDR) of 0.05. To find enriched annotations within hit genes, Gene Set Enrichment Analysis was performed for MC1- and MC1+ neurons using the Molecular Signatures Database C5 Ontology gene sets (Liberzon *et al*., 2011; Subramanian *et al*., 2005).

### Data availability

This paper does not report original code. Any additional information required to re-analyze the data reported in this paper is available from the lead contact upon request.

## ACKNOWLEDGEMENTS

We thank Dr. Marcos Otero-Garcia and Dr. Inma Cobos (Stanford) for the processed human AD scRNA-seq dataset; Dr. Manu Sharma (Weill Cornell) for sharing VAMP7^DN^ DNA constructs; Dr. Virginia Lee and University of Pennsylvania for the human post-mortem AD brains; Dr. Peter Davies for MC1 antibody; Dr. Rakez Kayed (UTMB) for TTC18 antibody; Dr. Xuming Tang and Dr. Chao Wang for technical support; Dr. Michael Ward (NIH), Dr. Wenjie Luo, Chinmay Vaidya, and Dr. Eileen Torres for discussions; Memorial Sloan Kettering Molecular Cytogenetics core for karyotyping; Weill Cornell Medicine Microscopy core for TEM; Weill Cornell Medicine Flow Cytometry core for FACS; Weill Cornell Medicine Genomics core for sequencing. This research was supported by the NIH U54NS100717, R01AG072758, R01AG054214, R01AG074541 (to L.G), and R01AG064239 (to W.L), Tau Consortium (to L.G. and M.K.) and JPB Foundation (to L.G.).

## AUTHOR CONTRIBUTIONS

L.G. and S.G. conceived the project. L.G., C.P.B, A.M.G., J.M.-P, Z.Z., A.S., S.G., and M.K. designed the experiments. C.P.B, A.M.G., J.M.-P, Z.Z., M.Y.W., A.E., L.F., M.M., M.Z., and S.G. performed the experiments. C.P.B, A.M.G., J.M-P., Z.Z., A.S., M.Y.W., L.F., T. Pozner., P.Y., T. Patel., A.Y., G.C., S.A., S.-A.M., W.L., V.M.Y.L., M.Z., M.K., and S.G. developed experimental protocols, tools, and reagents or analyzed data. C.P.B. and L.G. wrote the manuscript.

## Supplementary Figures

**Figure S1.**
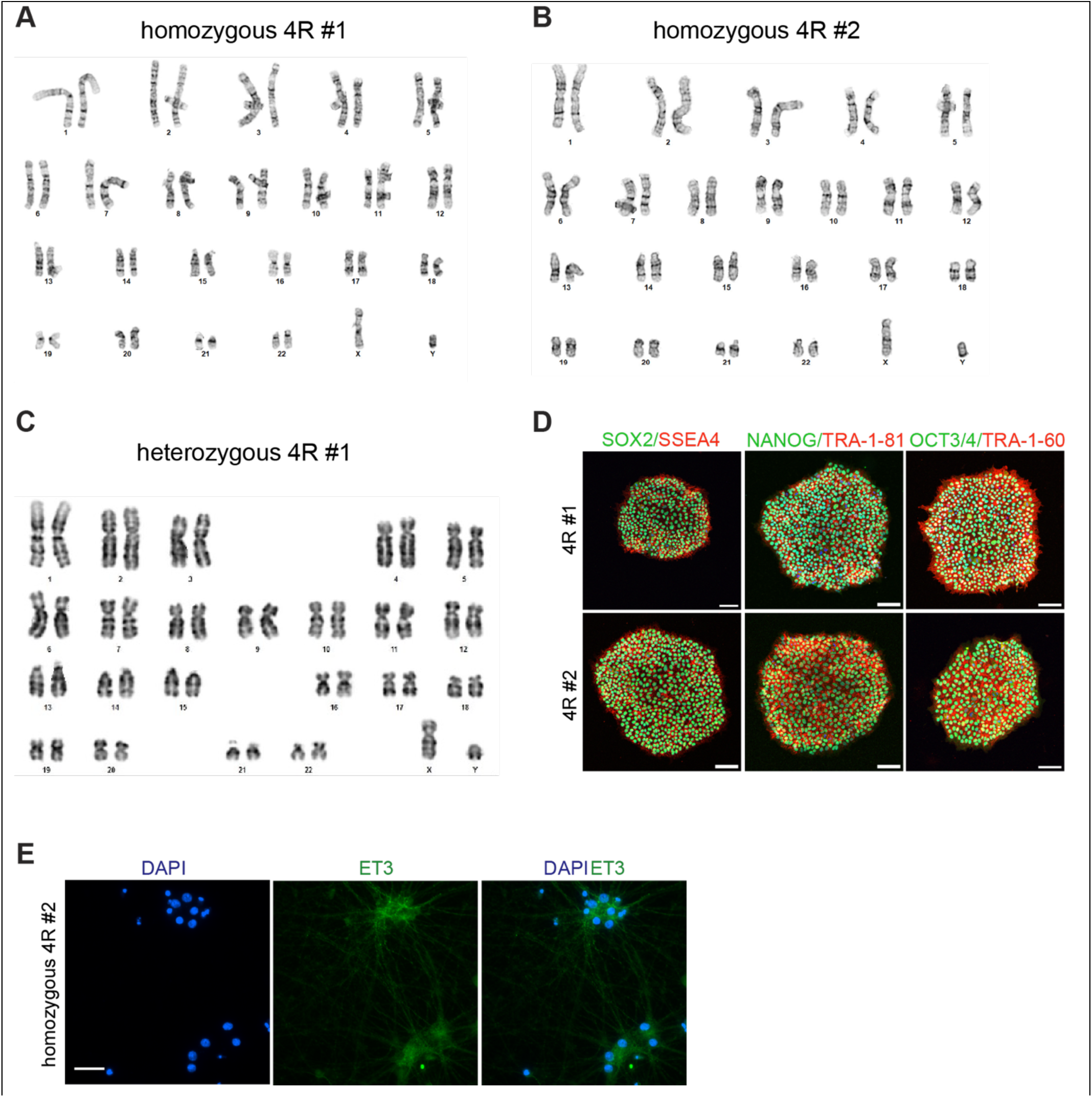
Generation of isogenic 4R-tau hiPSCs, related to Figure 1. (A and B) Normal karyotypes were confirmed for the 4R line clones #1 (A) and #2 (B). (C) A normal karyotype was confirmed for 95% of heterozygous 4R cells analyzed (clone #1). (D) Pluripotency marker staining (OCT4, TRA-1-60, NANOG, TRA-1-81, SOX2, SSEA4) confirms 4R (clone #1 and #2) hiPSC pluripotency. Scale bar, 100 μm. (E) Representative immunofluorescence images of 6-week-old (D44) i^3^N and 4R homozygous clone #2 neurons stained with ET3 and DAPI. Scale bar, 100 μm.

**Figure S2.**
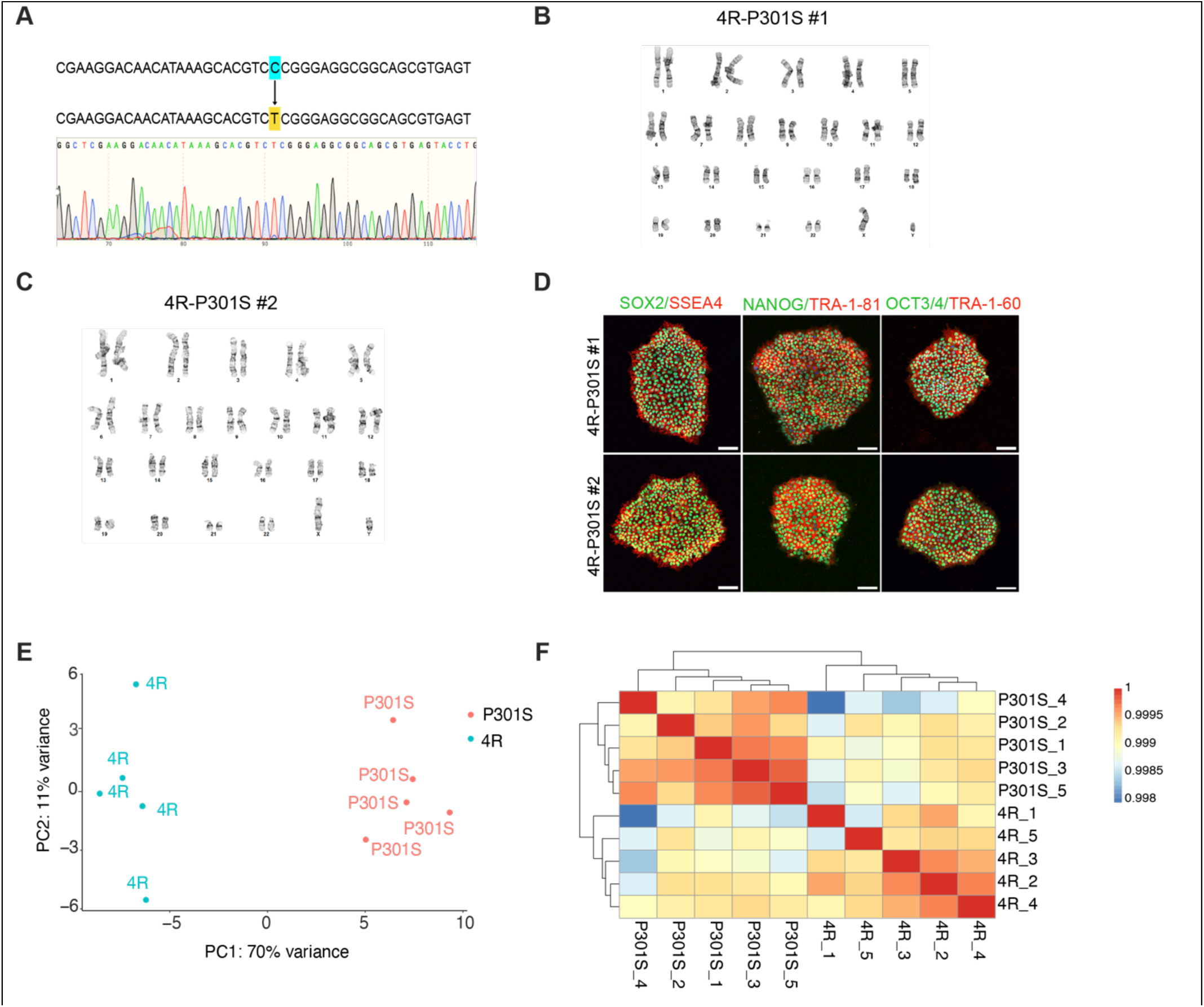
Generation and bulk RNA-sequencing of 4R-P301S hiPSCs, related to Figure 1. (A) Point mutation (C>T) validation was performed to generate the P301S mutation. (B and C) Normal karyotypes were confirmed for the 4R-P301S line clones #1 (B) and #2 (C). (D) Pluripotency marker staining (OCT4, TRA-1-60, NANOG, TRA-1-81, SOX2, SSEA4) confirms 4R-P301S (clones #1 and #2) hiPSC pluripotency. Scale bar, 100 μm. (E and F) Principal component analysis (E) and sample-sample correlation (F) of D43 4R and 4R-P301S neurons from bulk RNA-seq. n= five samples 4R, five samples 4R-P301S.

**Figure S3.**
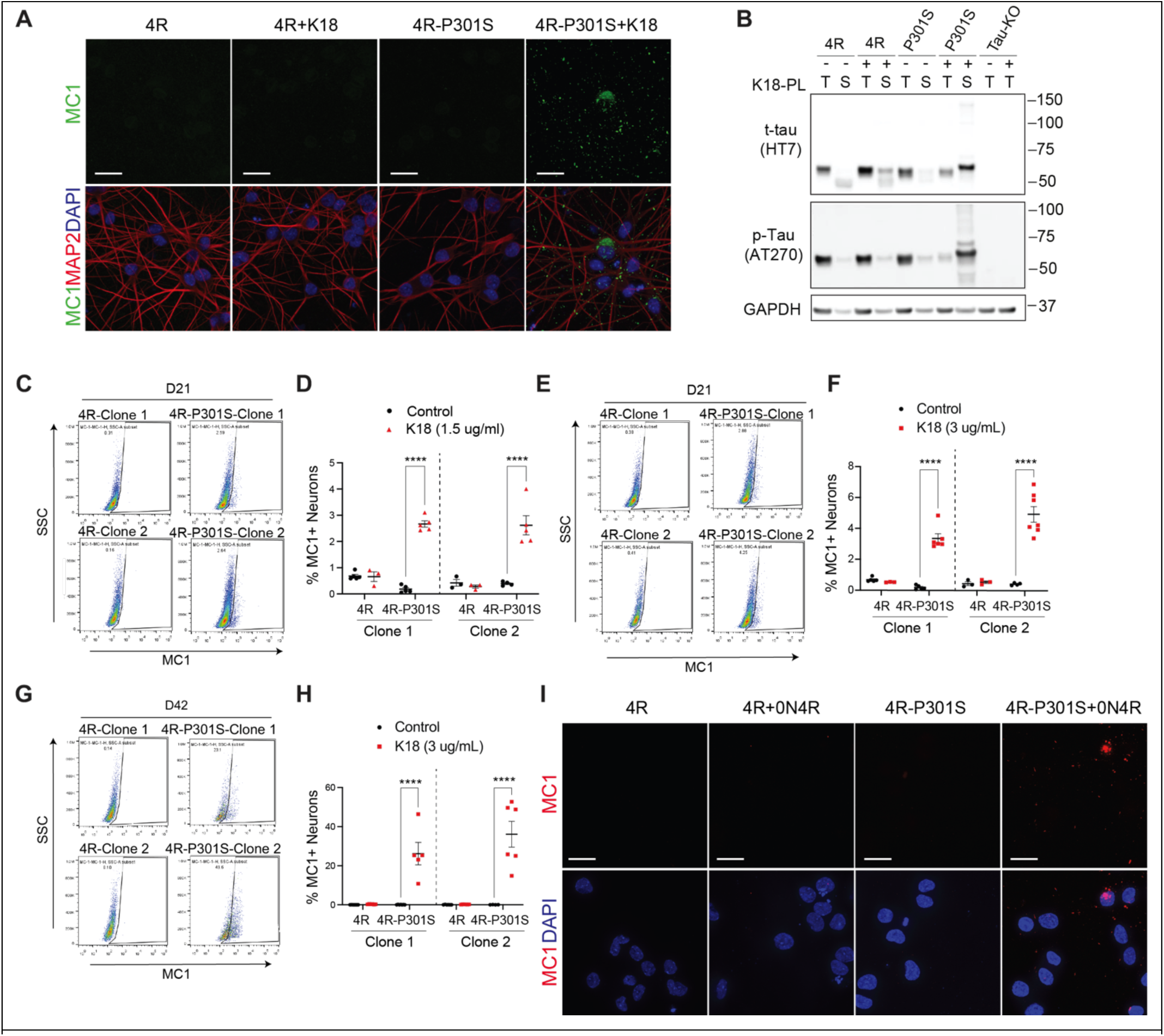
Modeling 4R-Tau inclusions, related to Figure 2. (A) Representative immunofluorescence images of D25(7+18) 4R and 4R-P301S neurons -/+K18 stained with MC1, MAP2, and DAPI. Scale bar, 25 μm. n=3, three independent experiments. (B) Immunoblot of phosphorylated tau (AT270) and total tau (HT7) from detergent-fractionated D68(22+46) 4R, 4R-P301S, and Tau-KO neuronal lysates. (C and D) Representative flow cytometry analysis (C) and quantification (D) of the percentage of MC1+ cells in D21(7+14) 4R and 4R-P301S neurons clone #1 and clone #2 -/+ 1.5 µg/mL K18 groups. n=3–6 biological replicates. **** p < 0.0001, two-way ANOVA, Šídák’s multiple comparisons test. (E and F) Representative flow cytometry analysis (E) and the quantification (F) of the percentage of MC1+ cells in D21(7+14) 4R and 4R-P301S neurons clone #1 and clone #2 -/+ 3 µg/mL K18 groups. n=3–6 biological replicates. **** p < 0.0001, two-way ANOVA, Šídák’s multiple comparisons test. (G and H) Representative flow cytometry analysis (G) and the quantification (H) of the percentage of MC1+ cells in D42(7+35) 4R and 4R-P301S neurons clone #1 and clone #2 -/+ 3 µg/mL K18 groups. n=3–6 biological replicates. **** p < 0.0001, two-way ANOVA, Šídák’s multiple comparisons test. (I) Representative immunofluorescence images of D25(7+18) 4R and 4R-P301S neurons -/+0N4R stained with MC1 and DAPI. Scale bar, 25 μm. n=4, three independent experiments.

**Figure S4.**
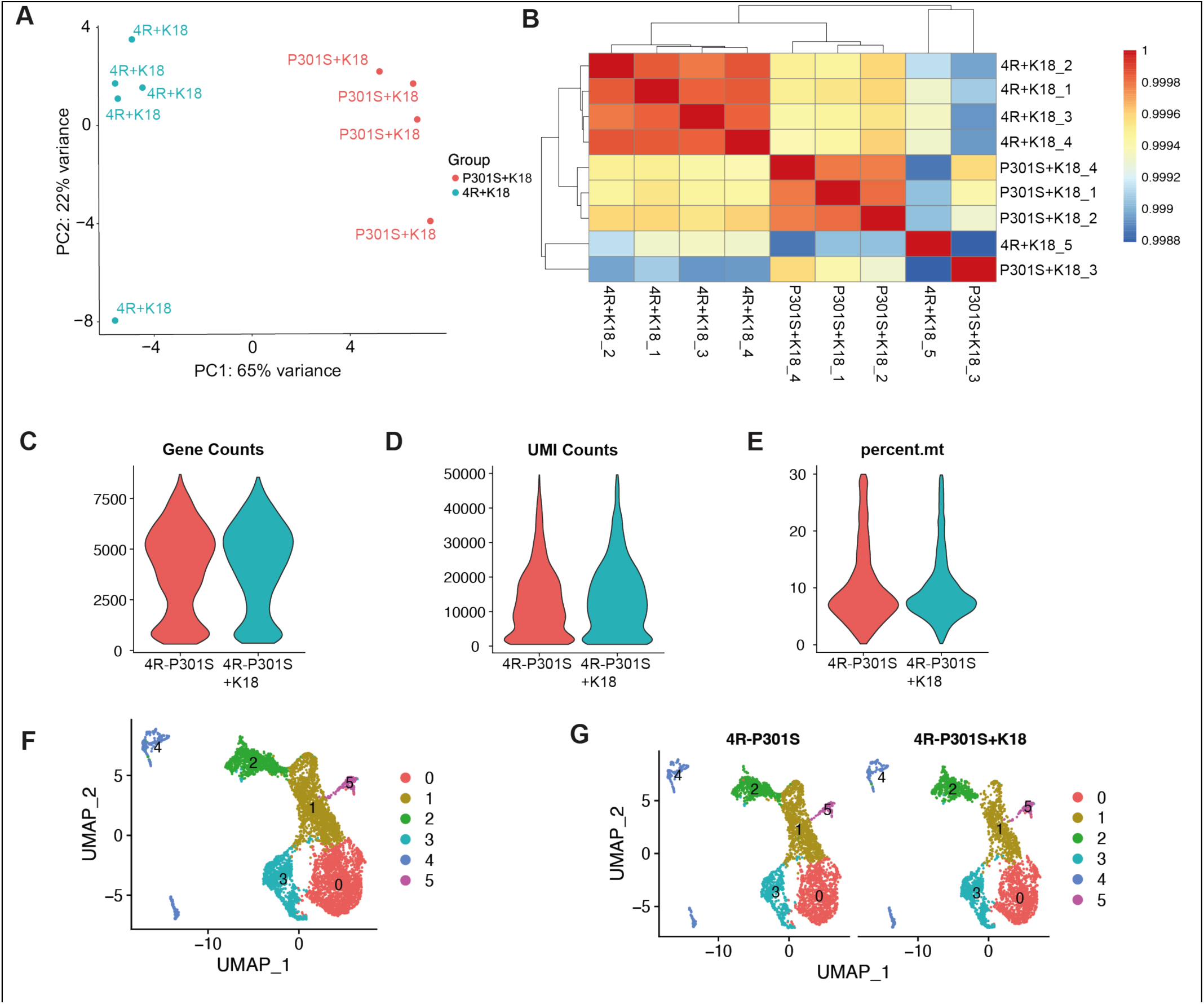
Bulk and single-cell transcriptomic analyses of 4R tauopathy model, related to Figure 2. (A and B) Principal component analysis (A) and sample-sample correlation (B) of D43(8+35) 4R and 4R-P301S neurons -/+ 1.5 µg/ml K18 from bulk RNA-sequencing. n=5 samples 4R+K18, 4 samples 4R-P301S+K18. (C-E) Violin plots showing spread of total genes (C), total UMIs (D), and percent of mitochondrial genes (E) detected per cell for each sample from single-cell RNA-sequencing of D39(7+32) 4R and 4R-P301S neurons. n=one sample P301S, one sample P301S+K18. (F and G) UMAP plot of all single cells (F) and separated by sample (G).

**Figure S5.**
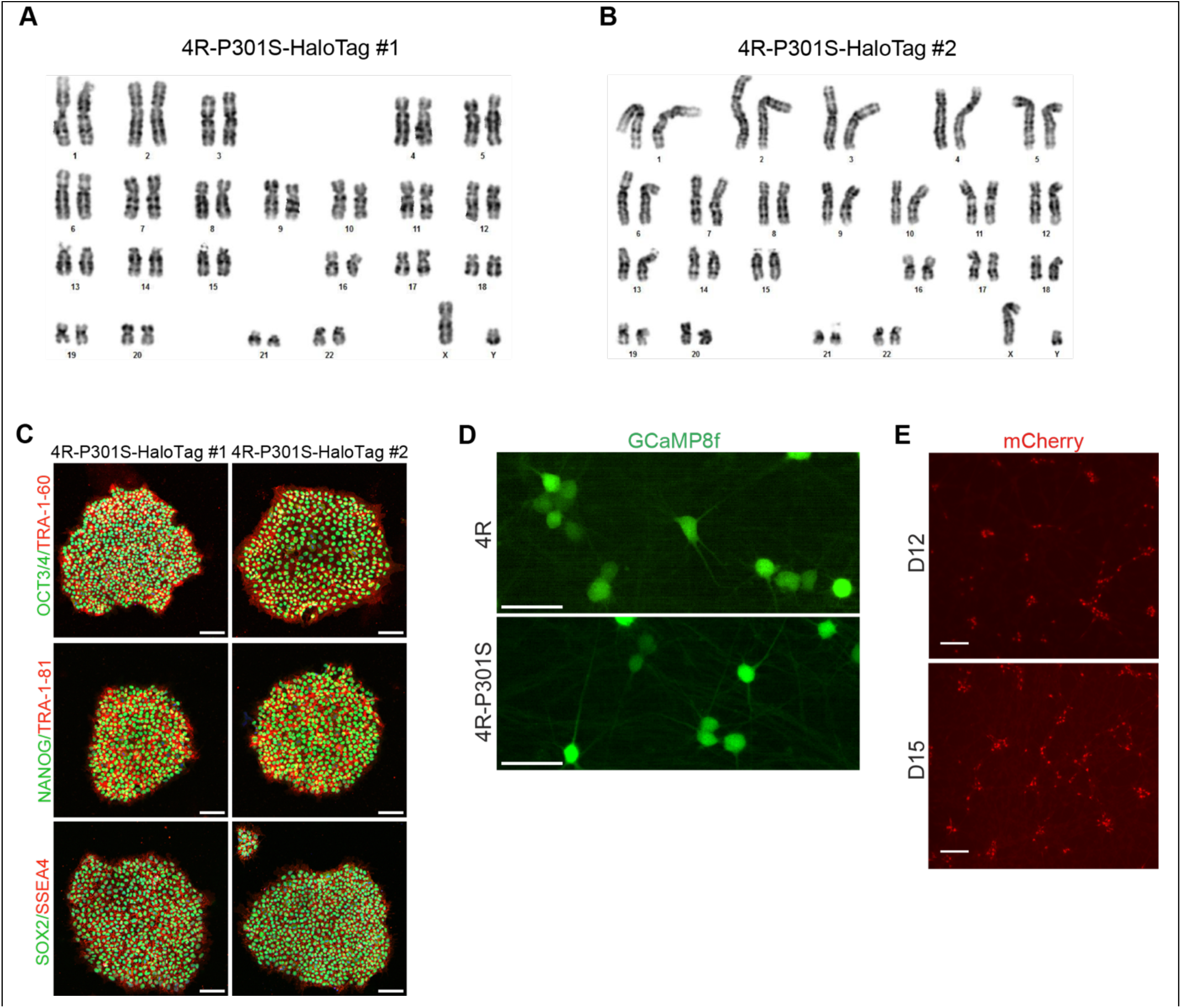
Generation of isogenic 4R-P301S-HaloTag hiPSCs and representative images of gCaMP8f- and hM4Di-transduced neurons, related to Figure 4. (A and B) Normal karyotypes were confirmed for the 4R-P301S-HaloTag line clones #1 (A) and #2 (B). (C) Pluripotency marker staining (OCT4, TRA-1-60, NANOG, TRA-1-81, SOX2, SSEA4) confirms 4R-P301S-HaloTag (clone #1 and #2) hiPSC pluripotency. Scale bar, 100 μm. (D) Representative image of D27 4R and 4R-P301S neurons overexpressing GCaMP8-fast genetically encoded calcium sensor at 488 nm, 30% light climate. Scale bar, 50 μm. (E) Representative fluorescent images of hM4Di-mCherry expression levels in live 4R-P301S-HaloTag neurons before (D12) and after (D14) CNO addition. Scale bar, 100 um.

**Figure S6.**
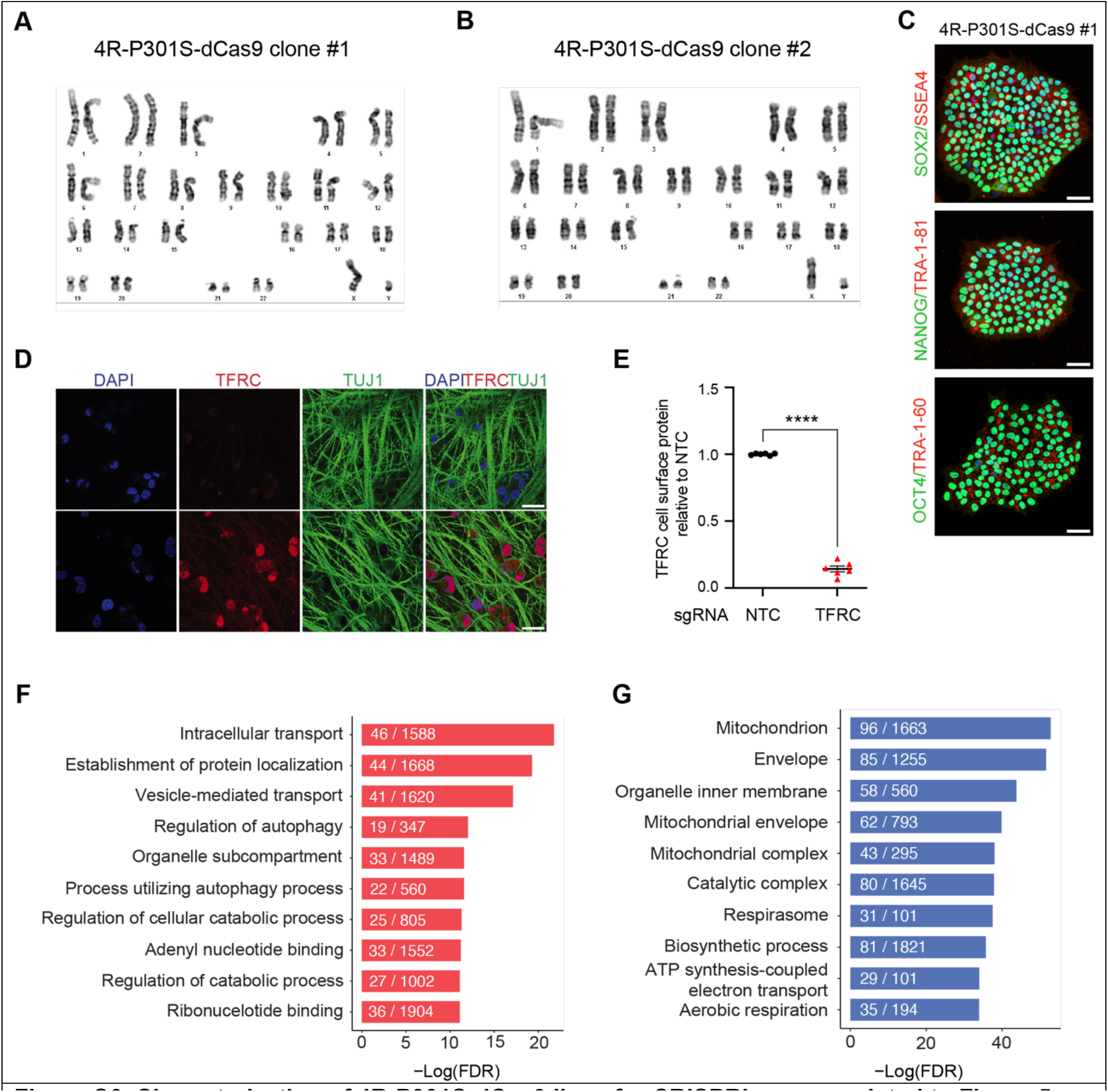
Characterization of 4R-P301S-dCas9 lines for CRISPRi screen, related to Figure 5. (A and B) Normal karyotypes were confirmed for the 4R-P301S-dCas9 line clone #1 (A) and clone #2 (B). (C) Pluripotency marker staining (OCT4, TRA-1-60, NANOG, TRA-1-81, SOX2, SSEA4) confirms 4R-P301S-dCas9 (clone #1) hiPSC pluripotency. Scale bar, 50 μm. (D) Representative immunofluorescence images of D21 4R-P301S-dCas9 neurons expressing a TFRC-targeting sgRNA or an NTC sgRNA stained with DAPI, TFRC, and TUJ1. Scale bar, 25 μm. (E) Functional validation of constitutively active CRISPRi activity by flow cytometry of TFRC surface protein level stained D21 4R-P301S-dCas9 neurons expressing a TFRC-targeting sgRNA or an NTC sgRNA. n=3, two independent experiments, ****p < 0.0001, unpaired t test. (F and G) Gene set enrichment analysis results for differentially expressed genes in MC1+ neurons (F) and MC1-neurons (G). Significantly enriched Gene Ontology terms for biological process, cellular component, and molecular function are shown.

**Figure S7.**
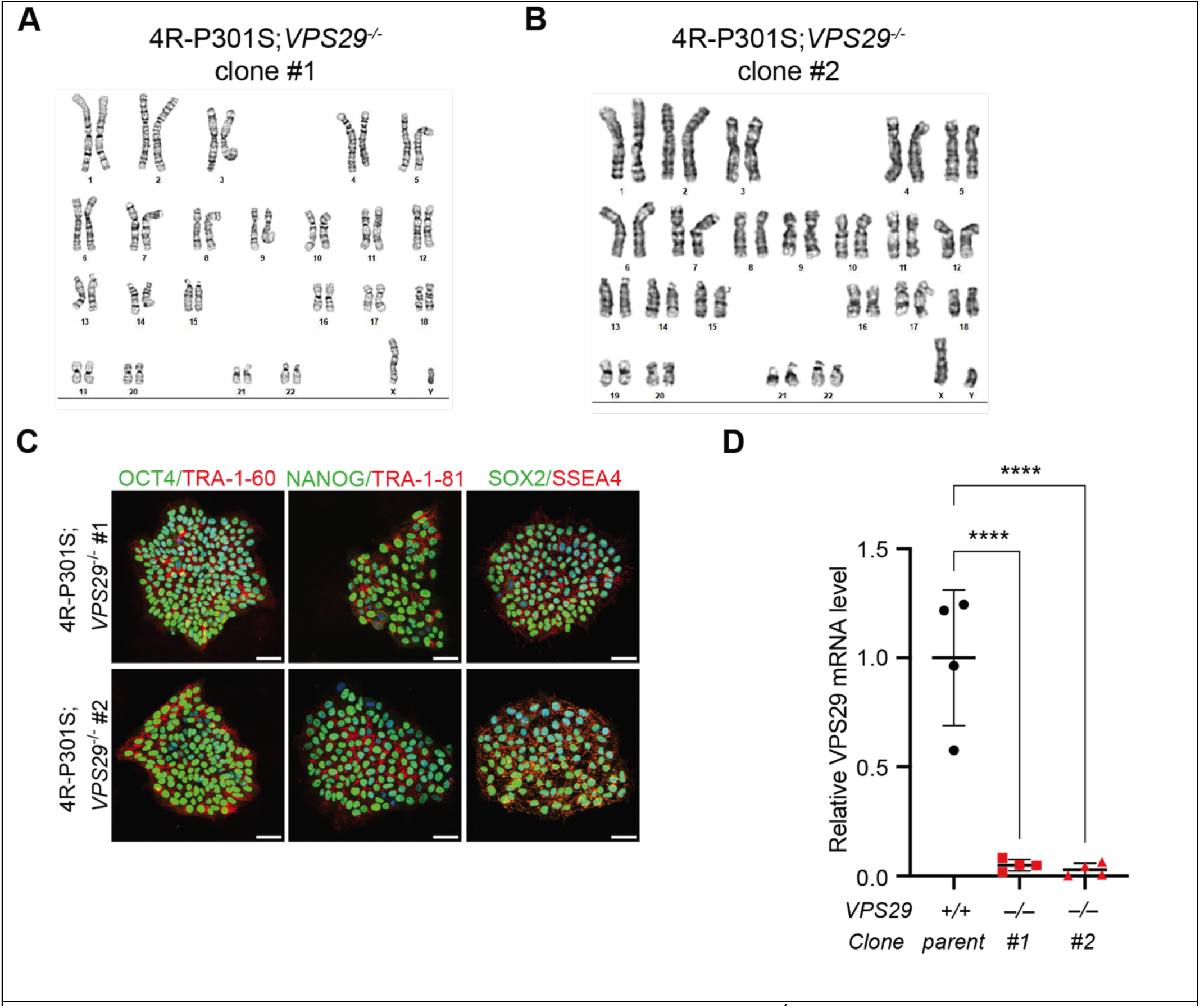
Characterization and validation of 4R-P301S;*VPS29^-/-^* hiPSCs, related to Figure 5. (A and B) Normal karyotypes were confirmed for the 4R-P301S;*VPS29^-/-^*line clone #1 (A) and in 95% of cells analyzed of clone #2 (B). (C) Pluripotency marker staining (OCT4, TRA-1-60, NANOG, TRA-1-81, SOX2, SSEA4) confirms 4R-P301S;*VPS29^-/-^*(clone #1, #2) hiPSC pluripotency. Scale bar, 50 μm. (D) Quantification of relative mRNA levels for *VPS29* in D21(7+14) K18-seeded 4R-P301S;*VPS29^-/-^*neuron clones #1 and #2 compared to parent. n=2 replicates, two independent experiments. **** p < 0.0001, one-way ANOVA and Dunnett’s multiple comparisons test.

**Figure S8.**
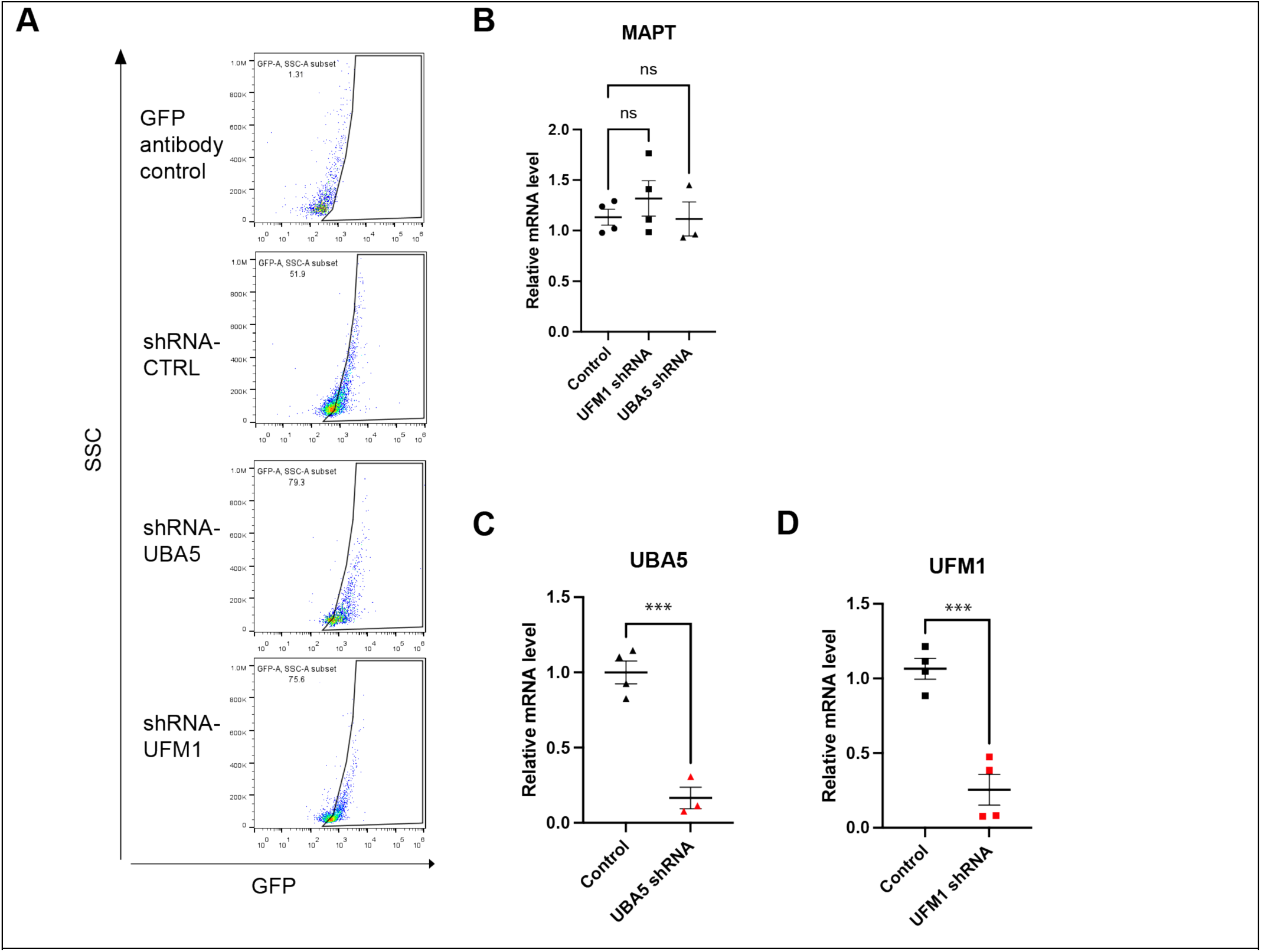
Validation of shRNA targeting UBA5 and UFM1 in 4R-P301S neurons, related to Figure 6. (A) Flow cytometry gating strategy for GFP+ neurons. (B-D) Quantification of relative mRNA levels for *MAPT* (B), *UBA5* (C), and *UFM1* (D) in D21(7+14) K18-seeded 4R-P301S neurons transduced with lentivirus containing shRNAs targeting *UBA5* and *UFM1*, compared to control shRNA. n=2 replicates, two independent experiments. *** p > 0.001, oneway ANOVA and Dunnett’s multiple comparisons test (B). unpaired t-test (C-D).

